# Spatial synchrony in vegetation response

**DOI:** 10.1101/520296

**Authors:** Haim Weissman, Yaron Michael, Nadav M. Shnerb

## Abstract

Spatial synchrony is ubiquitous in nature, and its decrease with the distance is an important feature that affects the viability of spatially structured populations. Here we present an empirical study of spatial synchrony in terrestrial vegetation using large scale remote sensing data. The decrease of synchrony with distance, as expressed by the correlation in rate of abundance change at a given time lag, is characterized using a power-law function with stretched-exponential cutoff. The range of these correlations appears to decrease when precipitation increases and to increase over time. The relevance of these results to the viability of populations is discussed.

## I. INTRODUCTION

Environmental variations are one of the main drivers of natural populations dynamics [13]. While the effect of demographic stochasticity diminishes with the size of a population, the relative strength of environmentally induced fluctuations is independent of population size. Accordingly, for medium-to-large size populations environmental stochasticity is expected to be the main cause of abundance fluctuations. A few recent large-scale empirical studies show that this is indeed the case [4, 10, 11, 14].

Under purely demographic noise, the time to extinction of a local population grows exponentially with its carrying capacity [12, 13, 18]. For reasonable parameters this usually yields unrealistically long persistence times, measured in thousands or millions of generations. The most probable path to extinction for local populations involves the cumulative effect of environmental variations that drive the population to very low densities. Under environmental fluctuations the time to extinction grows like a power of the carrying capacity, and when environmental stochasticity is relatively strong the exponent of this power is tiny, so the relevant extinction times are small [13, 22].

On spatial domains, the viability of a population reflects the balance between two processes: local extinction and migration. Migration acts to reduce the risk of extinction (rescue effect [3]) and allows for recolonization when a local population goes extinct. Therefore, spatial population synchrony [15], i.e., coincident changes of abundance (or biomass) in geographically disjoint populations, poses a threat of regional or global extinction, as nearby populations reach the brink of extinction together and cannot rescue each other.

Here we would like to report the level of synchrony observed for terrestrial vegetation systems, with particular emphasis on the arid and semi-arid climatic zones. Taking advantage of the wide spatial coverage and high spatial resolution of remotely sensed data, we have calculated the spatial correlations in the ratios of abundance [6] with 10 × 10 meters resolution.

The Normalized Difference Vegetation Index (NDVI) has been used as a measure of local biomass. This technique cannot distinguish between different vegetation species, so the measured correlations reflect the synchronous fluctuations of the biomass. These correlations most likely caused by a common response of different plant species to changes in environmental factors such as temperature and precipitation. A very similar technique was implemented recently by Defriez and Reuman [5] to identify global patterns of synchrony, for completely different length scales (their spatial resolution is 1°).

It turns out that the spatial correlations of the response decay slowly over short length scales, and cross over to rapid decay above some typical distance *λ* (see Figure 2 below). We discovered that a correlation function in which a power-law decay is truncated by a stretched exponential cutoff is flexible enough to yield very good fits to our data, and used this function

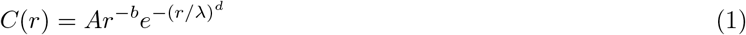

to characterize the results.

Spatial synchrony was measured for each rain line in our data, with resolution of 50mm/year. When we have analyzed the ratios of abundance in successive years, we discovered that the typical cutoff scale decays when the precipitation level increases, where in some extreme cases the power-law region shrinks to zero and the decay of synchrony is purely stretched exponential. Similar results were obtained for 2 years lag. Using another dataset (with resolution of 30 × 30 meters, so the results are more noisy) we examined spatial synchrony in 13 years lag. The stretched exponential truncated power law still provides nice fits, but the typical cutoff scale *λ* is much **longer**, where in some cases the results seem to indicate a pure power-law over the measurement region.

## II. METHODS

### A. the data

We have used consecutive satellite images of the Sahel region in North Africa and of the Australian desert (Fig 1). As a measure of biomass density the most commonly used [7] Normalized Difference Vegetation Index was implemented. To produce NDVI layers, we used data from Sentinel 2 where each pixel (our spatial resolution) corresponds to an area of 10 × 10 meters. To exclude images with high cloud cover, we applied the greenest pixel composite filter [19], in which the pixel that has the highest green reflectivity value was selected from the images available within the date range. Areas were stratified with (average) rainfall lines (taken from [8]).

**FIG. 1:**
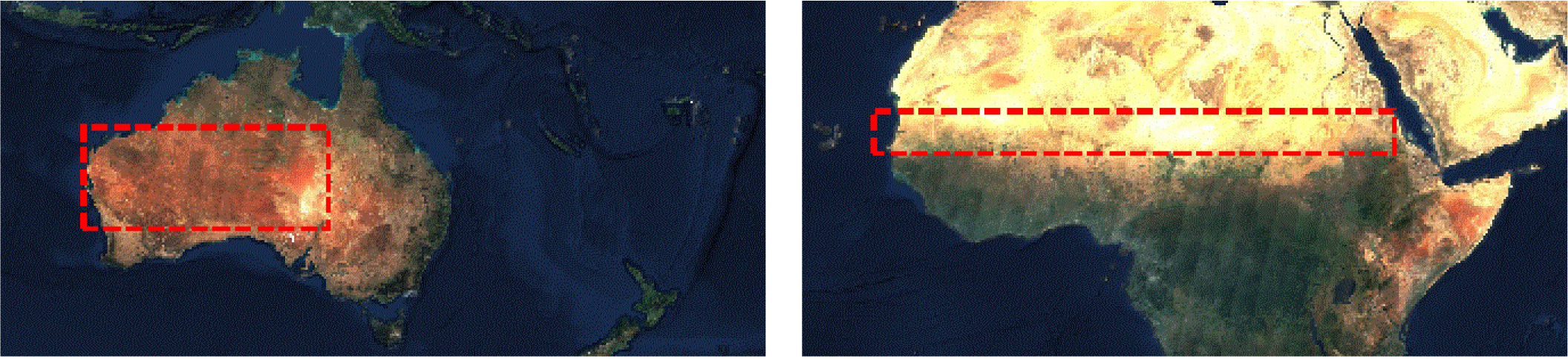
Our study areas contain the Sahel region in Africa (latitude: 11.7° − 15.1° and longitude: 17.8° − 40°, total area 3 × 10^6^ km^2^, right panel) and the desert of Australia (latitude: 20.0° − 30.0° and longitude: 113.0° − 140.0°, total area 2.5 × 10^6^ km^2^, left panel). Satellite images were taken from Sentinel-2 cloudless (https://s2maps.eu) by EOX IT Services GmbH (Contains modified Copernicus Sentinel data 2016 and 2017).

Data from the Sahel was recorded between 15/09/2016 – 15/10/2016 and 15/09/2017 – 15/10/2017, and from Australia we compared records taken between 01/04/2016 – 31/05/2016, 01/04/2017 – 31/05/2017 and between 01/04/2018 – 31/05/2018, so for Australia we can report results for one year lag (T=1) and two yeas lag (*T* = 2). For *T* = 13 we used the Sahel data (EVI index) taken in 2002 and 2015, with 30 × 30 meters resolution. A complete description of these datasets was presented in [20].

### B. Local response and its correlations

For each pixel we have two NDVI indices (for the same pixel in two different years) that we regard as proxies for the local biomass density *ρ*. The relative growth or decline of the biomass in that pixel is defined as,

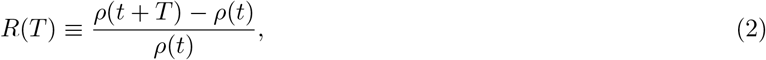

We then measured the two dimensional autocorrelation of *R* values as a function of distance *r*. To do that, for each distance *r* between 10m and 20km we sampled at least 10^7^ pairs of pixels whose distance from each other is *r*. For each *r* each measurement yields two numbers *R*_1_ and *R*_2_, and the corresponding correlation is,

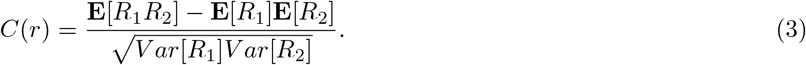

## III. RESULTS

### A. The Sahel

A typical result is shown in Figure 2, where the correlation *C*(*r*) is depicted against *r* for the African Sahel region, 400-450 rainline. To obtain values of *b*, *λ* and *d* we fitted the expression (1) to the logarithmicaly binned data using a double logarithmic scale, but (as shown in the insets) the same parameters yield a decent fit for the row data on both double logarithmic scale and on arithmetic scale.

**FIG. 2:**
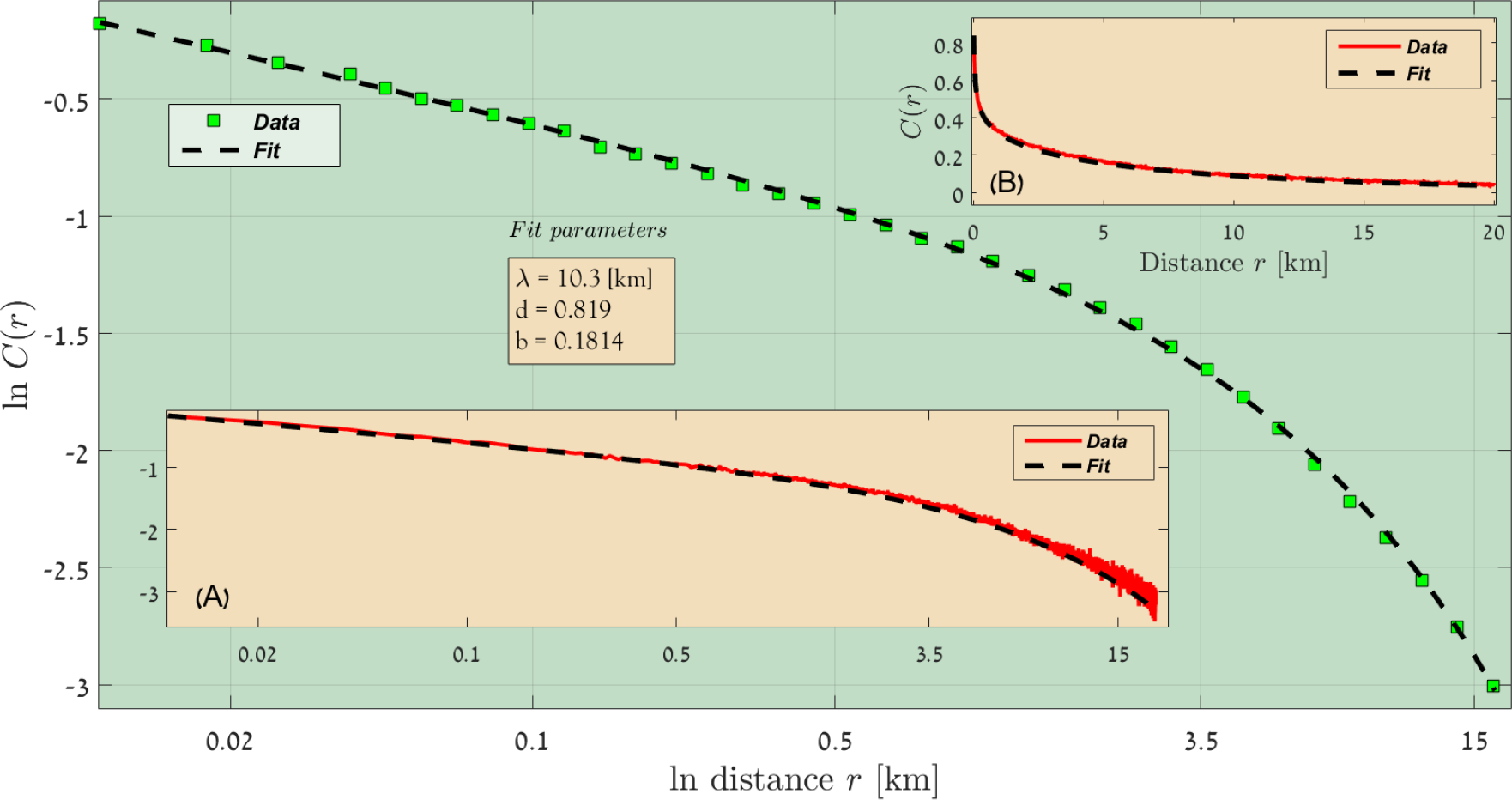
*C*(*r*) vs. *r* for the 400 – 450mm/year region of the Sahel (double logarithmic scale). *C*(*r*) was calculated for all distances between 10 meters and 20 kilometers, the results were logarithmically binned and are presented in the main panel (green squares). This dataset was then fitted to the stretched-exponentially truncated power law (Eq. 1), and the best fit parameters *b*; *λ* and *d* are given in the mid inset. The dashed black line corresponds to Eq. (1) with these parameters and actually fits nicely the data. In the insets we present the calculated correlations (before logarithmic binning) on a linear (B) and double logarithmic (A) scale (full red line), together with the graph of Eq. (1) with the *same* parameters, and one can see that the quality of the fit does not depend on the binning or on the type of scale used.

All other results from the Sahel are given in Supplementary A. One sees that the stretched-exponential truncated power-law is indeed flexible enough to fit all these results. Note that we do not pretend to argue that Eq. (1) is *the* law that describes the decay of spatial synchrony in vegetation systems. Instead, these nice fits allow the use of three parameters: *b*, *λ* and *d* - in order to characterize the behavior of the empirical lines.

Qualitatively, *b* measures the short-distance rate of decline in synchrony. The smaller is *b*, the slower is that decay. *λ* is the characteristic distance above which the spatial synchrony starts to decay rapidly, and *d*, the shape parameter, measures how rapid is the decay above *λ*. When *d* = 1 the decay is exponential, while *d* < 1 implies softer decay.

Figure 3 shows how these three parameters change with the level of precipitation. The parameter *b* characterizes the short-range decay. It takes values between 0.25 and zero, and decreases as precipitation increases, so the short-range decay slows down. On the other hand the crossover from slow to fast decay takes place at *λ*, and this scale factor also becomes smaller when the rain increases. Finally the shape factor *d* decreases along rainlines, so the decay becomes softer and softer.

**FIG. 3:**
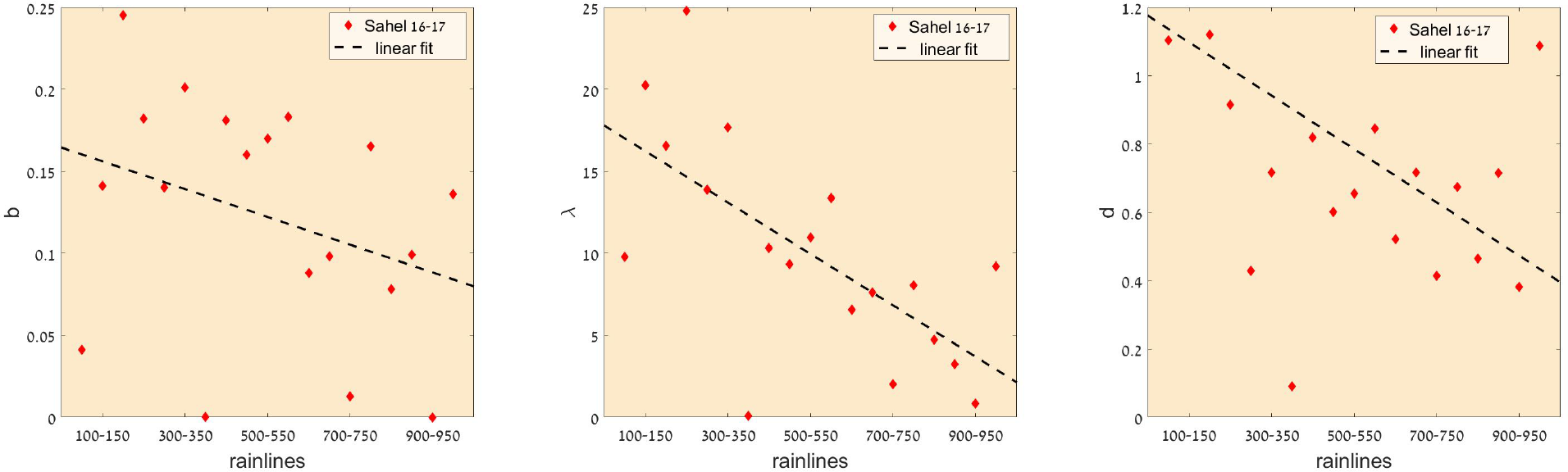
The variations of the short distance power-law *b* (left), the scale parameter *λ* (middle) and the shape parameter *d* (right) as obtained for the one year lag Sahel data. As precipitation increases, the value of all parameters decreases, as demonstrated by the linear fit to these results, depicted by a dashed black line. The exceptional point *d* = 2.6 for the 100 – 150mm/year region is not seen in the right panel.

For the Sahel we can study another dataset that was already analyzed (in different context) in [20]. Here we compare the EVI index for 2002 and 2015 (*T* = 13), in 30 × 30 meters per pixel resolution. Although the data is much more noisy and the fits are less impressive, the characterization of the measured *C*(*r*) by the stretched-truncated power-law (1) is still good in most cases, as demonstrated in Supplementary B. The results for *T* = 13 are summarized in Fig 4. Interestingly, *C*(*r*) for *T* = 13 decays much more slowly that the spatial synchrony for *T* = 1. For *T* = 1 the scale factor *λ* is below 25 kilometers (with a few cases - e.g., the 350-400 rainline - where *λ* and *b* approach zero, so the decay is a pure stretched exponential without short-range power-law). On the other hand,the range of *λ* for *T* = 13 is around 100 kilometers, and in some cases the fit assigned even larger values to *λ*, so the result is essentially a pure power-law decay.

**FIG. 4:**
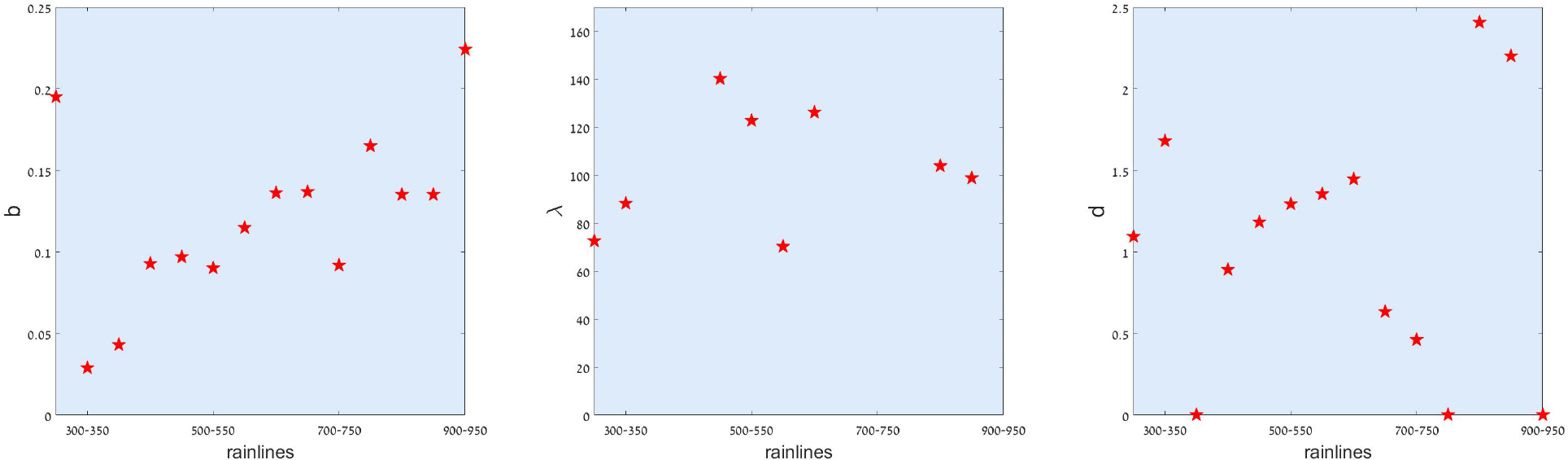
The variations of the short distance power-law *b* (left), the scale parameter *λ* (middle) and the shape parameter *d* (right) as obtained for the 13 year lag (2002-2015) Sahel data. The fitted values of *λ* for rainlines 350-450, 750-800 and above 900 are very large, meaning that there is essentially no exponential cutoff to the power-law as may be seen in the corresponding figures in Supplementary B. In these cases the value of *d* is also meaningless.

If spatial synchrony for *T* = 1 reflects the extent of spatially correlated climatic events (like the amount of rain during a specific wet season), and if these events are uncorrelated between years, then one should expect *λ* to decrease with *T*. The empirically observed increase seems to suggest that the long-time growth rate of vegetation is affected by different mechanism (for example, temperature variations) with much higher spatial correlation length. Another possibility is that the long-term correlations have to do with extreme rain events that have much larger spatial extent than typical rainstorms.

### B. Australian desert

For the Australian desert we have NDVI 10 × 10 meters resolution data from three consecutive years, 2016-2018. Accordingly, we present results for two *T* = 1 lag and one *T* = 2. All the graphes are presented in supplementary C, and the corresponding characteristic parameters are shown in Figure 5.

**FIG. 5:**
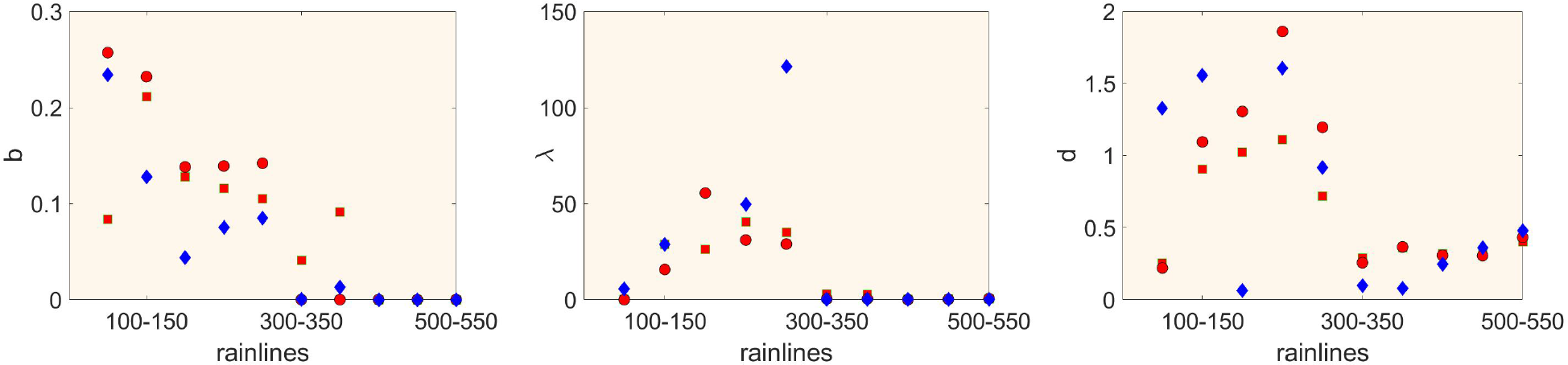
The variations of the short distance power-law *b* (left), the scale parameter *λ* (middle) and the shape parameter *d* (right) as obtained from the Australian desert data. Red circles correspond to the one year lag 2016-2017, red squares represent *T* = 1, 2017-2018 and blue diamonds correspond to *T* = 2, 2016-2018.

In Australia one observes a relatively sharp transition around the 300mm rainline. Above this line *λ* is very small, there is no power-law regime ane the decay of spatial synchrony is described by a pure stretched exponent. Below the 300mm/year line the spatial decay crosses over from short-distance power law to long distance stretched exponential behavior, as in the Sahel. There are no significant differences between the *T* = 1 and *T* = 2 datasets.

## IV. DISCUSION

In this paper we presented a preliminary attempt to describe the spatial synchrony in terrestrial vegetation systems using remotely sensed data. The pros and the cons are clear: on the one hand, the number of elementary pixels is huge (for each rainline we typically had more than a billion pixels) so the statistics is relatively good. On the other hand, the timeseries are short and the technique do not discriminate between different species or between trees and shrubs.

Spatial synchrony is usually attributed to some combination of three factors: it may reflect strong migration among local patches (in such a case the persistence time of a regional population decreases when the migration is too strong, yielding a bell-shaped persistence-migration relationship [23]), trophic interaction with other mobile populations (moving predators) or the autocorrelations of environmental fluctuations (Moran effect [17]). Among these, the *T* = 1 synchrony patterns reported here seem to be a manifestation of the Moran effect. One may speculate that the increase of spatial synchrony in the *T* = 13 may be related to the long-term effects of seed dispersal, but of course other mechanisms may yield the same effect as discussed above.

The decay of spatial synchrony with distance was captured by the correlation function *C*(*r*) and was characterized, by fitting the empirical results with Eq. (1), by three parameters: the power-law exponent *b*, the scale parameter *λ* and the shape parameter *d*. Our analysis suggest two general trends: faster spatial decay of *C*(*r*) as precipitation increases, and slower decay as the lag *T* grows.

The dynamics of spatial populations reflects the balance between local extinctions and recolonization by migrants (or seeds) from nearby patches. The populations persist if extinctions are compensated by colonization, otherwise it goes extinct. The simplest model that captures these features is the contact process, a lattice model with local extinction and nearest neighbor colonization. This model supports a continuous extinction transition with the typical exponents of the directed percolation equivalence class [9]. In the active phase the population reaches a finite density, in the inactive phase it decays exponentially to zero, and only at one point, the transition point, the decay is described by a power-law. Practically, almost all models of population dynamics on spatial domains show this type of extinction transition, as long as both extinctions and recolonizations are local [2, 21].

An important exception to this statement is the case when population is exposed to *global* temporal fluctuations [1]. In that case the transition is completely different, and a temporal Griffith phase (a finite parameter region in which the population density decays like a power law) emerges between the extinction phase and the active phase.

The result presented above suggest that the temporal environmental variations which affects vegetation dynamics are spatially correlated over relatively long distances. Although we did not show global fluctuations, up to few kilometers the correlations decay like a power-law with relatively small exponent, and the power-law region seems grow with time (so for *T* = 13 it sometimes reaches 20 kilometers). In closely related models [16], spatial correlations that decay so slowly were shown to affect strongly the characteristics of the transition, making it closer to the one observed for the global temporal fluctuations case. If a similar argument is valid for the vegetation system, the actual extinction transition (that may be related to important regime shifts like desertification) may differ substantially from the predictions of a naive theory that takes into account only local environmental fluctuations.

## Acknowledgements

We acknowledge Ryan Chisholm for helpful discussions and comments. We acknowledge the support of the Israel Science Foundation, grant no. 1427/15.

