## Supplementary material for "Spatial synchrony in vegetation response": Supp

**Supplementary materials for:**
**Spatial synchrony in vegetation response**

Haim Weissman[1], Yaron Michael[2] and Nadav M. Shnerb[1]
[1]*Department of Physics, Bar-Ilan University, Ramat-Gan IL52900, Israel.*
[2]*Department of Geography and Environment, Bar-Ilan University, Ramat-Gan IL52900, Israel.*

### Introduction

In this supplementary we present the correlations as measured in the Sahel and in the Australian desert (see Fig 1 of the main text), for $T = 1$, $T = 2$ and $T = 13$, for different rain lines between 50mm/year and 1000mm/year. Together with the correlations we present, in each figure, the fits to

$$C(r) = Ar^{-b}e^{-(r/\lambda)^d}, \tag{1}$$

i.e., to a stretched-exponentially truncated power law with the parameters $b$ (the exponent of the power law), $\lambda$ (the scale parameter) and $d$ (the shape parameter).

In all these figures we have used the same presentation technique that we have implemented in Fig. 2 of the main text, and $C(r)$ is plotted vs. $r$ for the relevant spatial region. $C(r)$ was calculated for all distances between 10 meters and 20 kilometers, the results were logarithmically binned and are presented in the main panel of each figure by green squares. This dataset was then fitted to the stretched-exponentially truncated power law (Eq. 1), and the best fit parameters $b$; $\lambda$ and $d$ are given in the mid inset of each figure. The dashed black line corresponds to Eq. (1) with these parameters. In the insets we present the calculated correlations (before logarithmic binning) on a linear (B) and double logarithmic (B) scale (full red line), together with the graph of Eq. (1) with the $b$; $\lambda$ and $d$ parameters listed in the mid-panel.

For more details see the Method section in the manuscript.

In section A, figures 1-19 show the results for the Sahel with $T = 1$ (2016-2017).

In section B, figures 20-36 show the results for the Sahel with $T = 13$ (2002-2015).

In section C.1, figures 37-46 show the results for the Australian desert with $T = 1$ (2016-2017).

In section C.2, figures 47-56 show the results for the Australian desert with $T = 1$ (2017-2018).

Finally, in section C.3 figures 57-66 show the results for the Australian desert with $T = 2$ (2016-2018).

### A. Results from Sahel, 1 year lag, 2016-2017

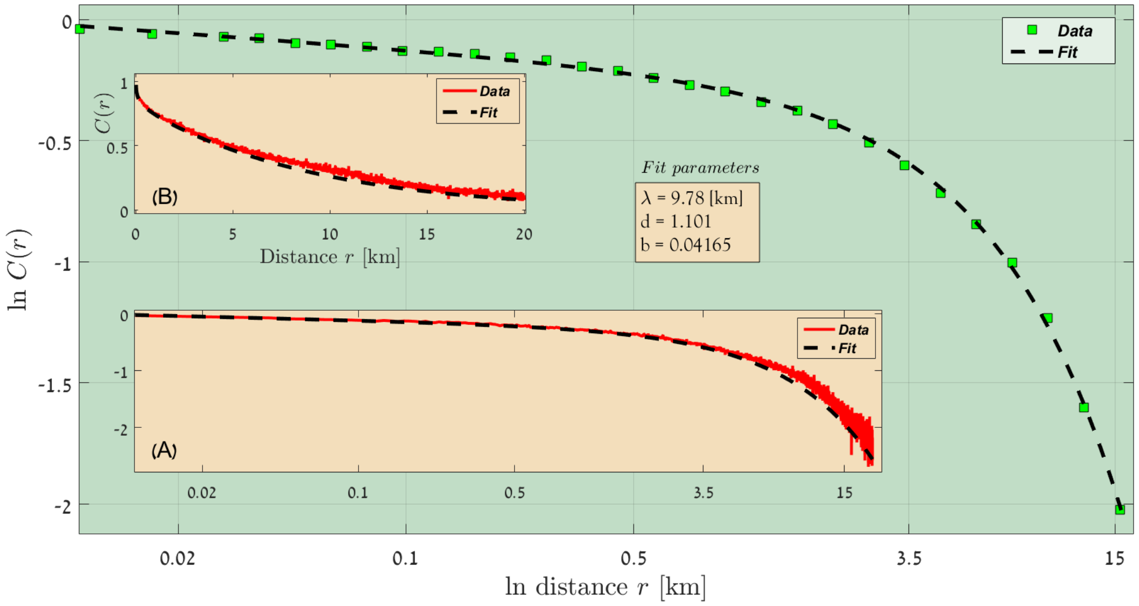


FIG. 1: Sahel, Rainfall lines 50 - 100 [mm/year], 2016-2017.

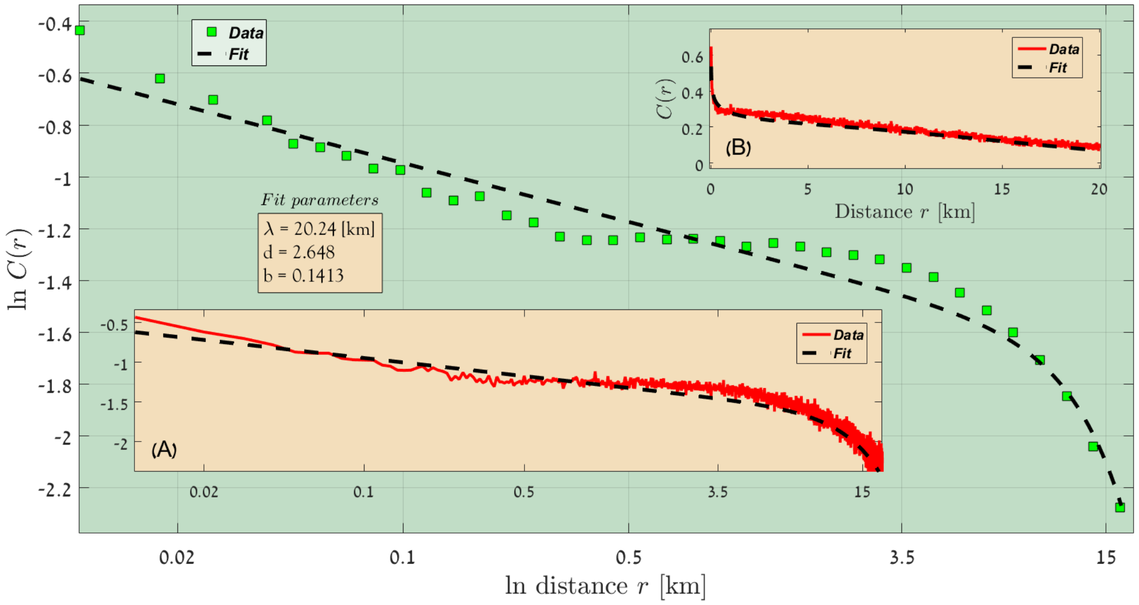


FIG. 2: Sahel, Rainfall lines 100 - 150 [mm/year], 2016-2017.

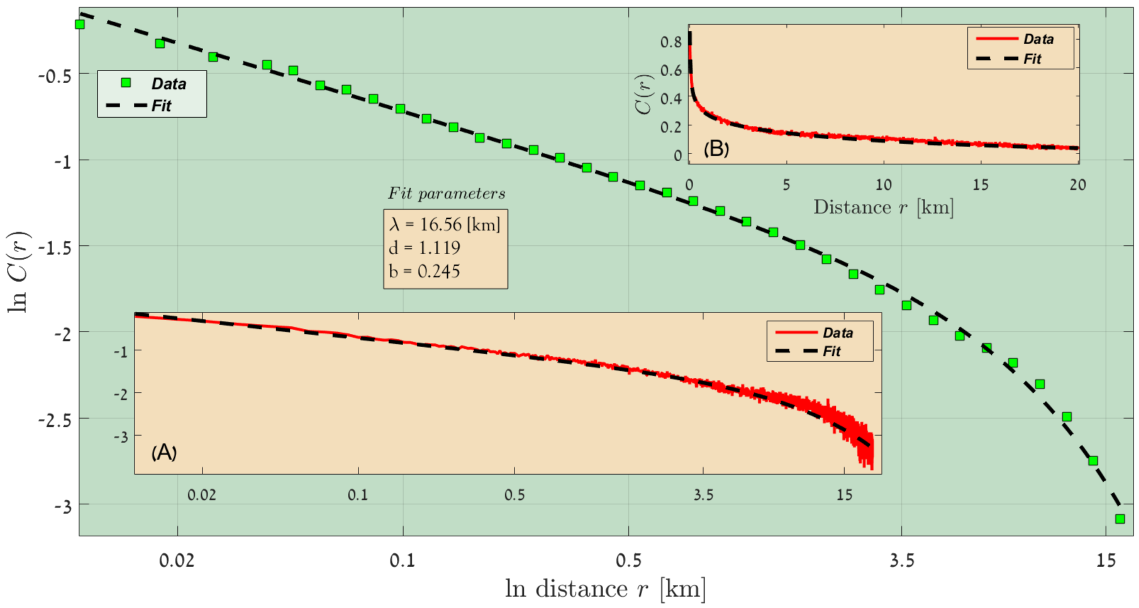


FIG. 3: Sahel, Rainfall lines 150 - 200 [mm/year], 2016-2017.

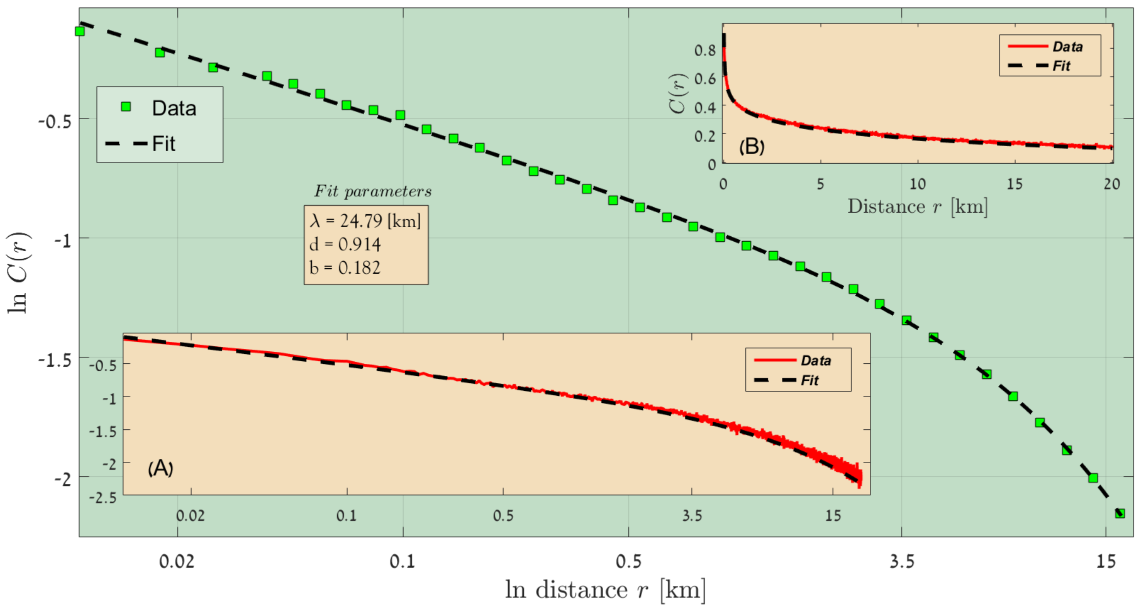


FIG. 4: Sahel, Rainfall lines 200 - 250 [mm/year], 2016-2017.

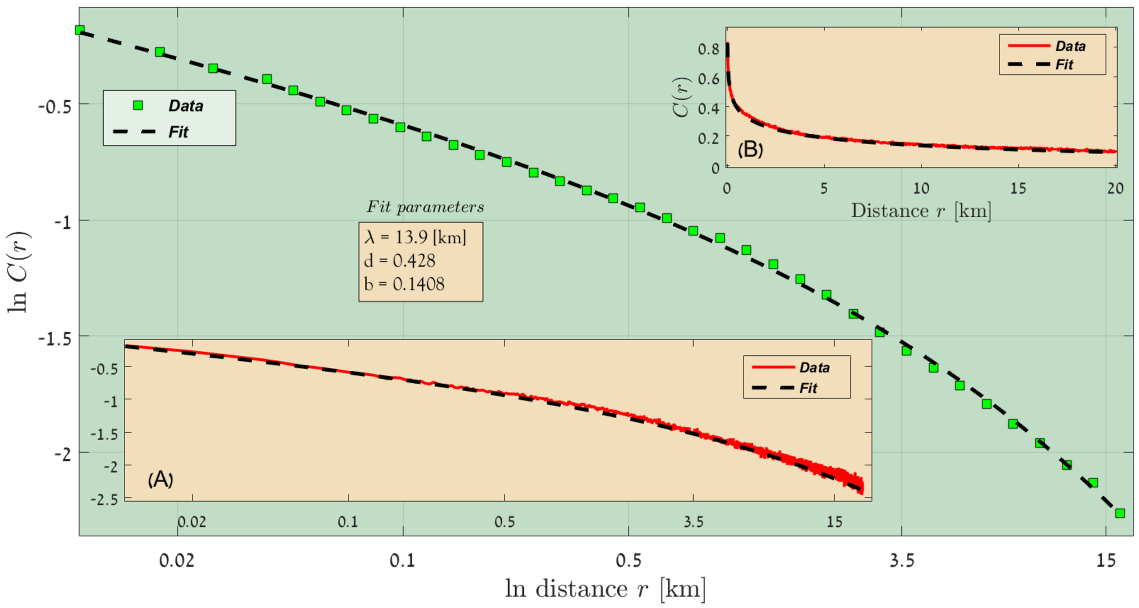


FIG. 5: Sahel, Rainfall lines 250 - 300 [mm/year], 2016-2017.

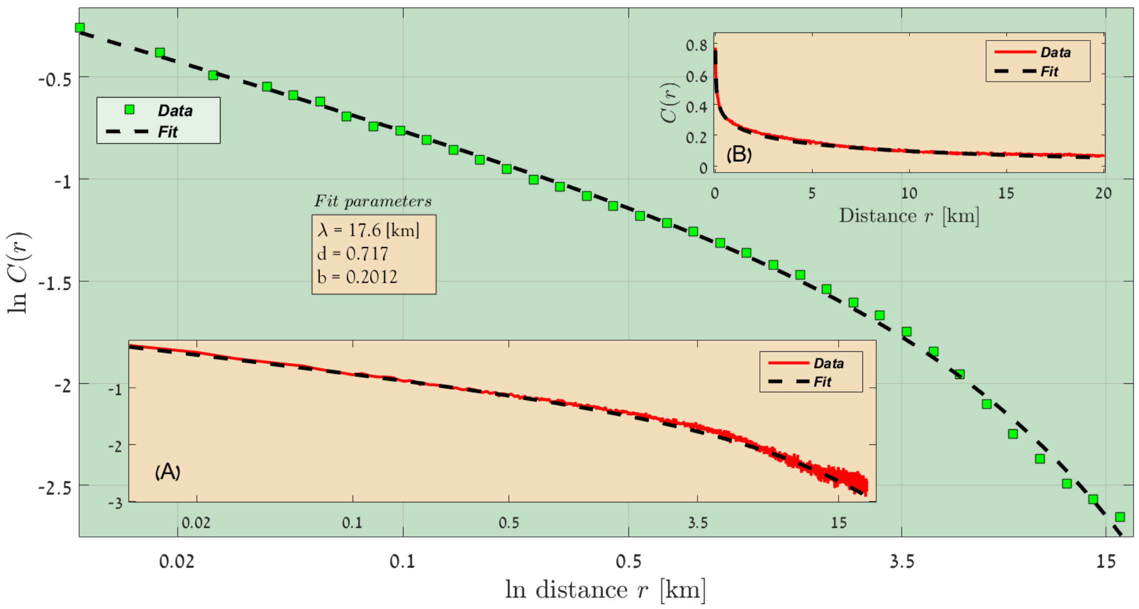


FIG. 6: Sahel, Rainfall lines 300 - 350 [mm/year], 2016-2017.

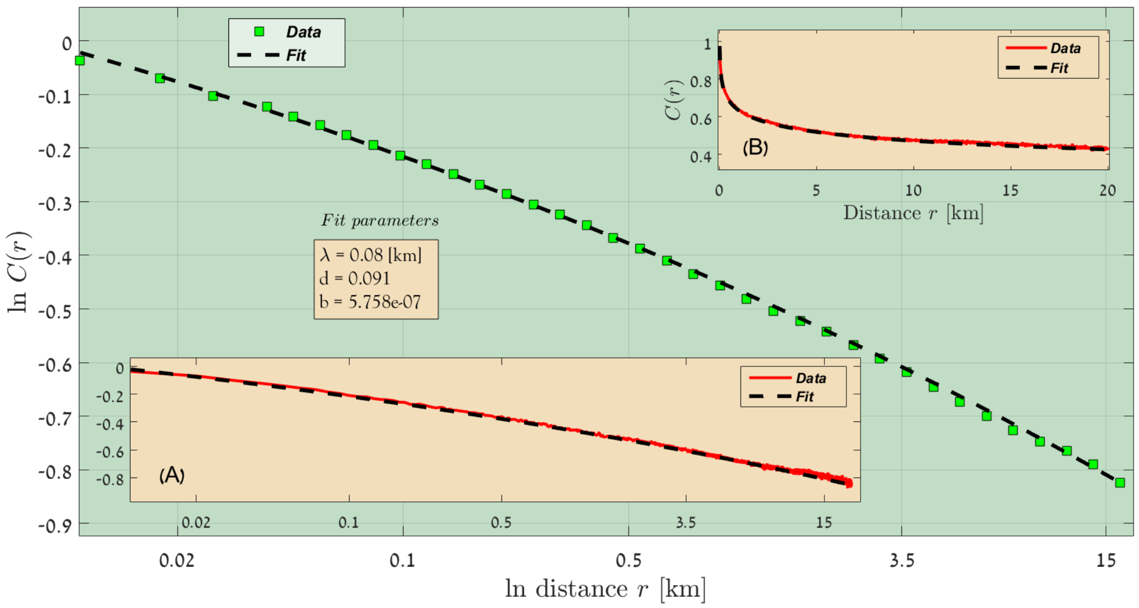


FIG. 7: Sahel, Rainfall lines 350 - 400 [mm/year], 2016-2017.

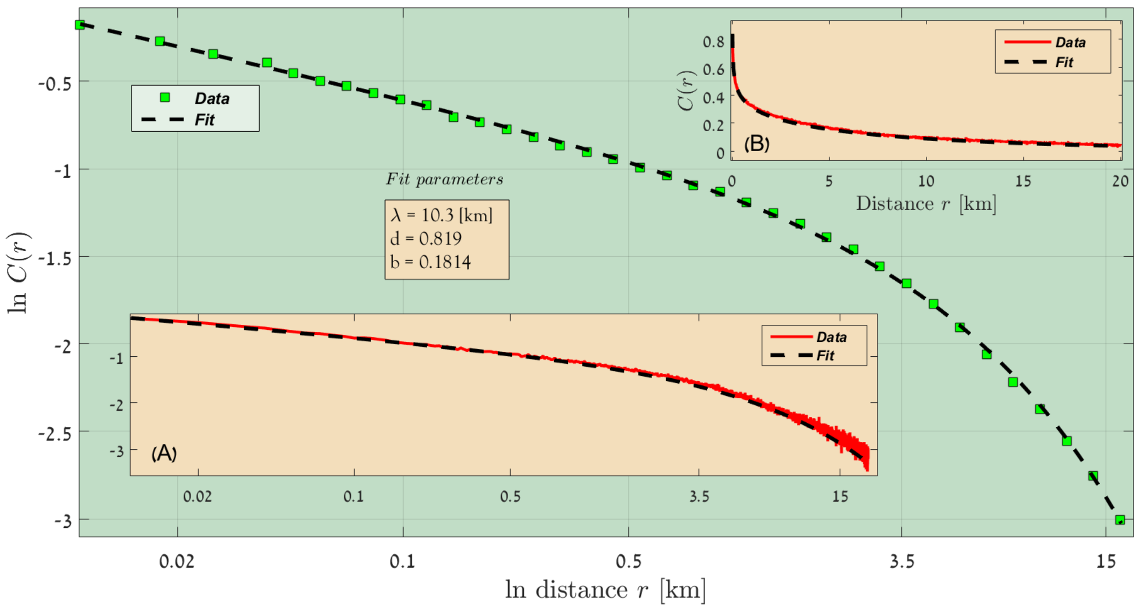


FIG. 8: Sahel, Rainfall lines 400 - 450 [mm/year], 2016-2017.

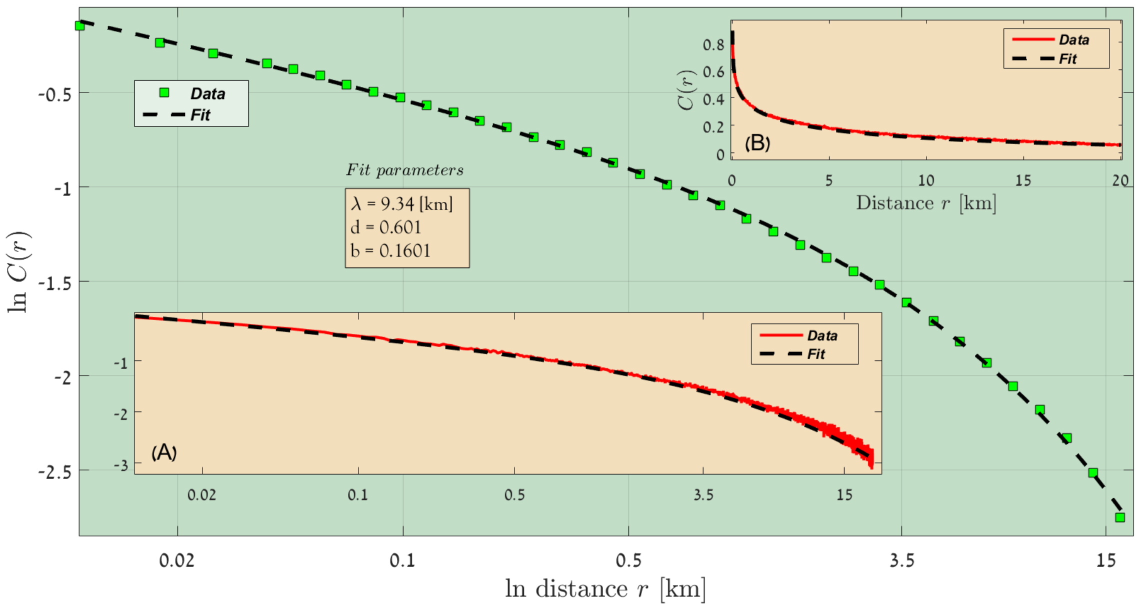


FIG. 9: Sahel, Rainfall lines 450 - 500 [mm/year], 2016-2017.

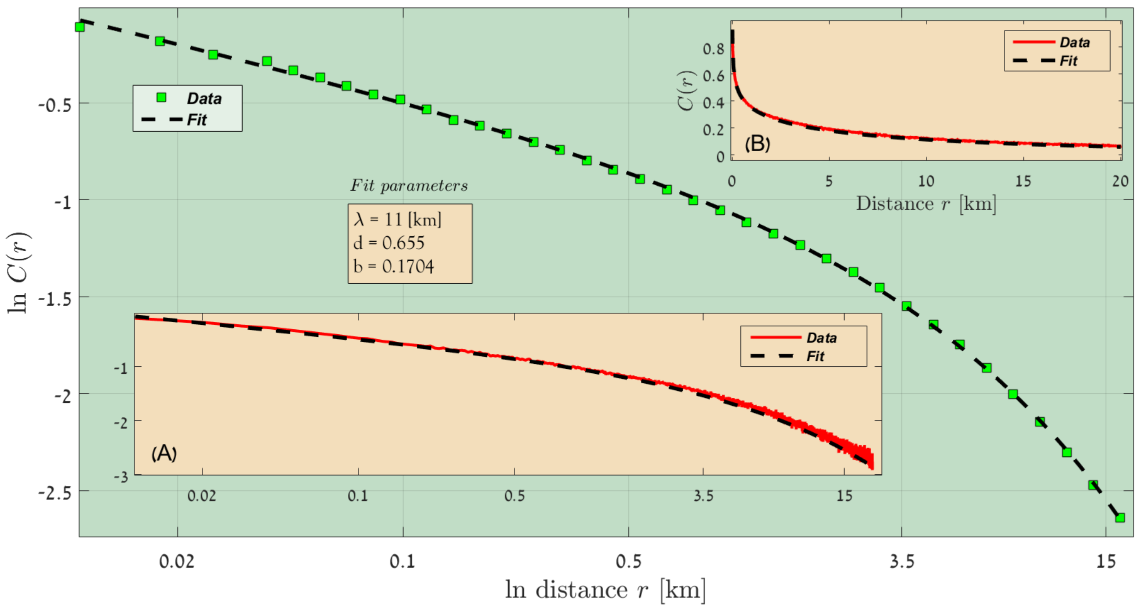


FIG. 10: Sahel, Rainfall lines 500 - 550 [mm/year], 2016-2017.

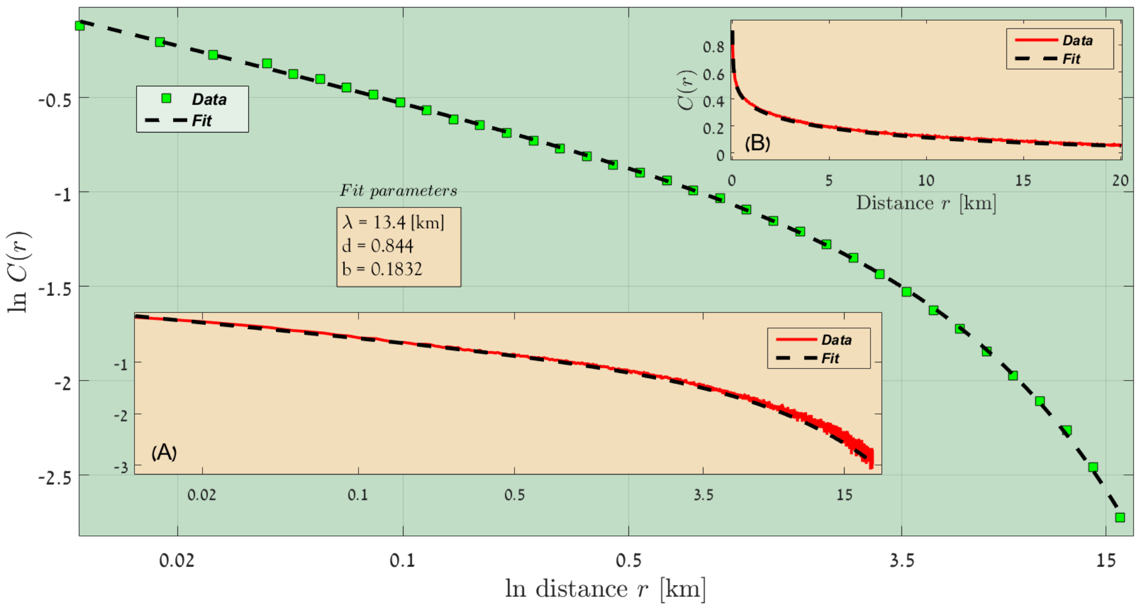


FIG. 11: Sahel, Rainfall lines 5500 - 600 [mm/year], 2016-2017.

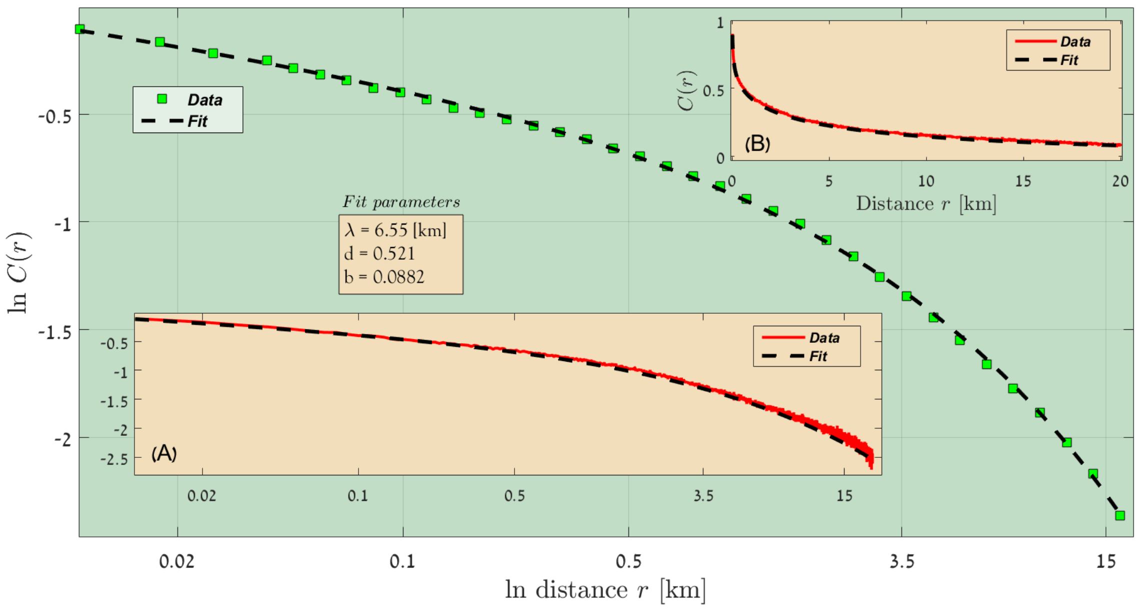


FIG. 12: Sahel, Rainfall lines 600 - 650 [mm/year], 2016-2017.

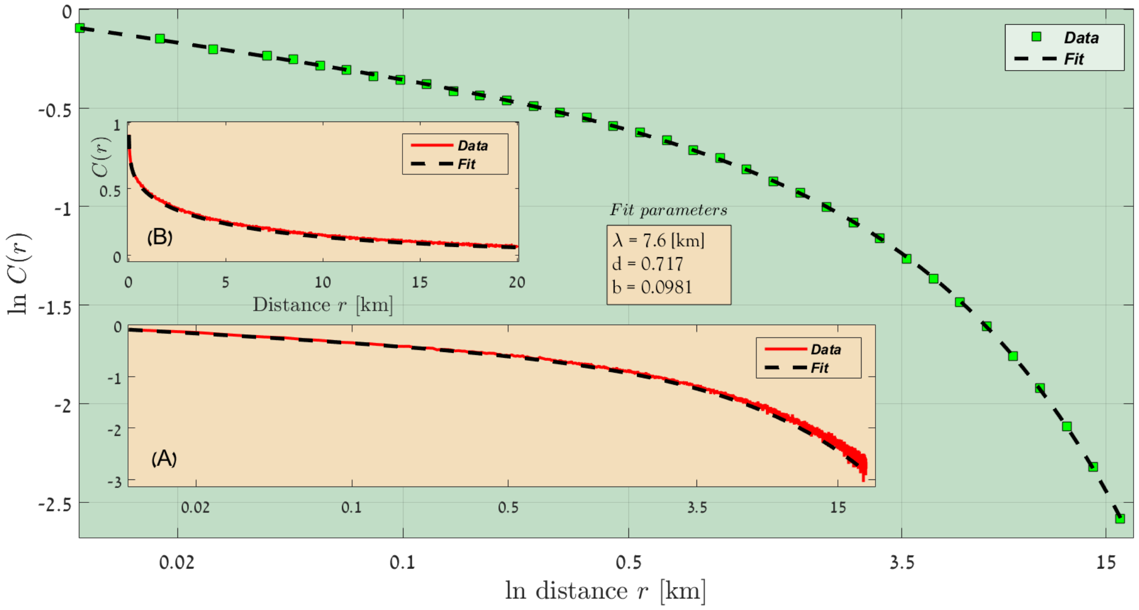


FIG. 13: Sahel, Rainfall lines 650 - 700 [mm/year], 2016-2017.

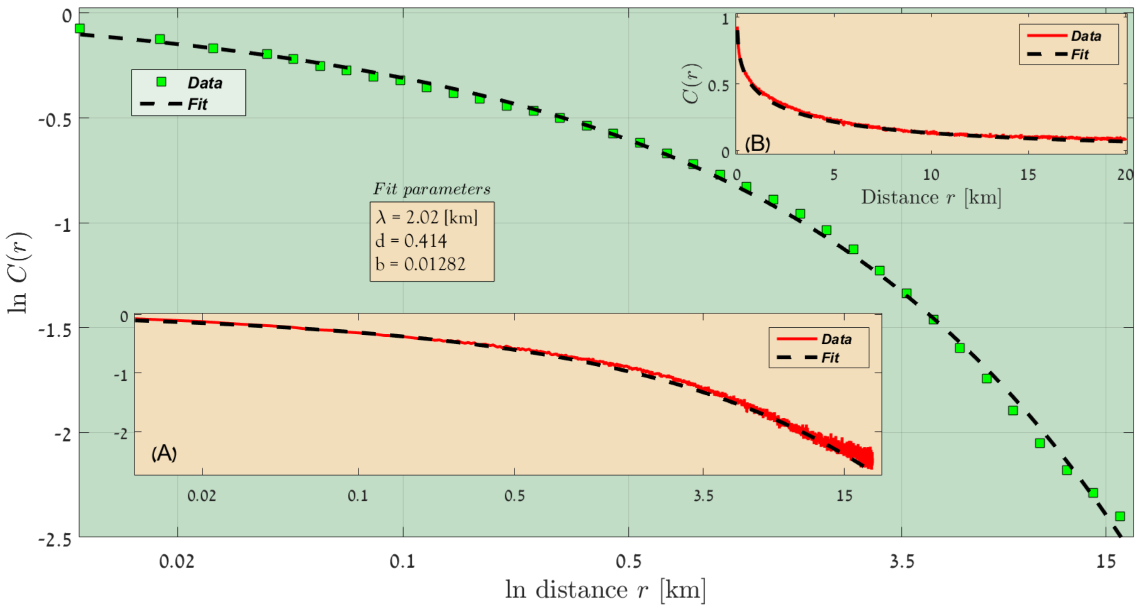


FIG. 14: Sahel, Rainfall lines 700 - 750 [mm/year], 2016-2017.

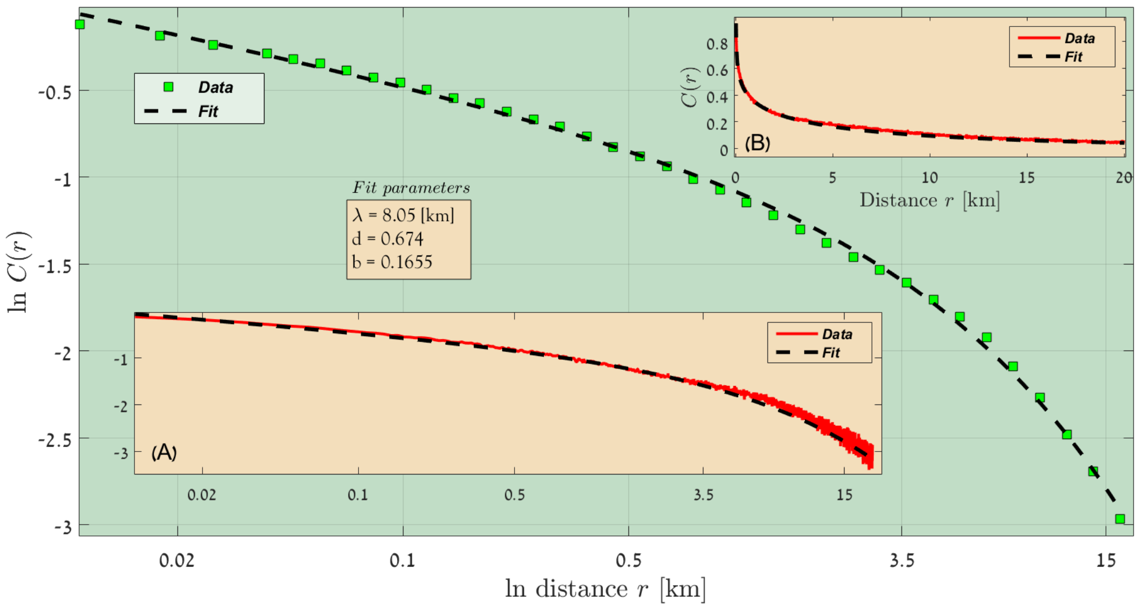


FIG. 15: Sahel, Rainfall lines 750 - 800 [mm/year], 2016-2017.

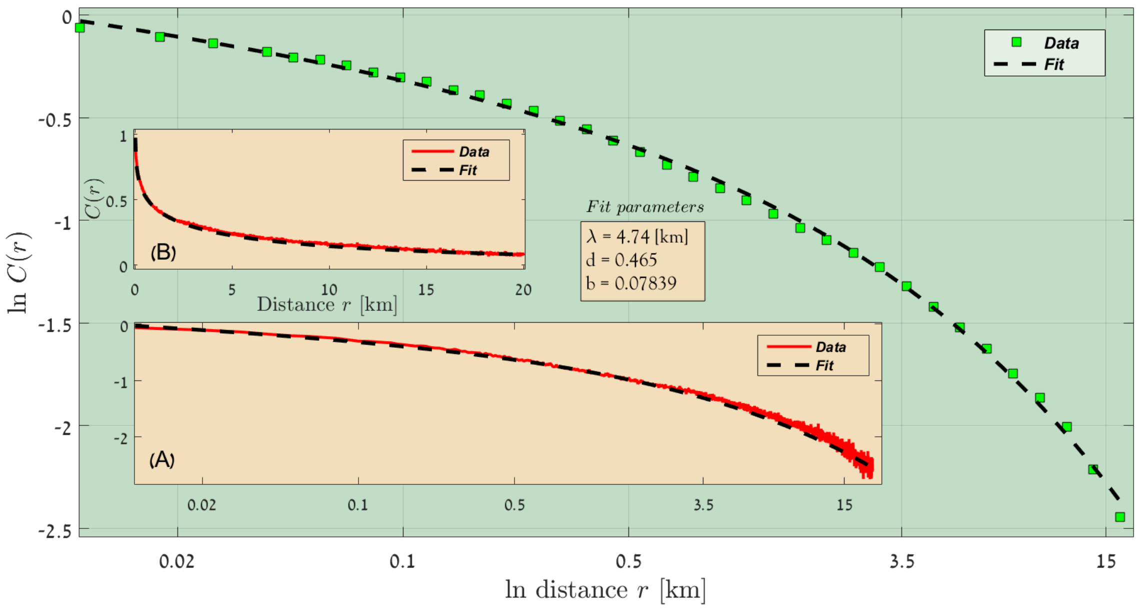


FIG. 16: Sahel, Rainfall lines 800 - 850 [mm/year], 2016-2017.

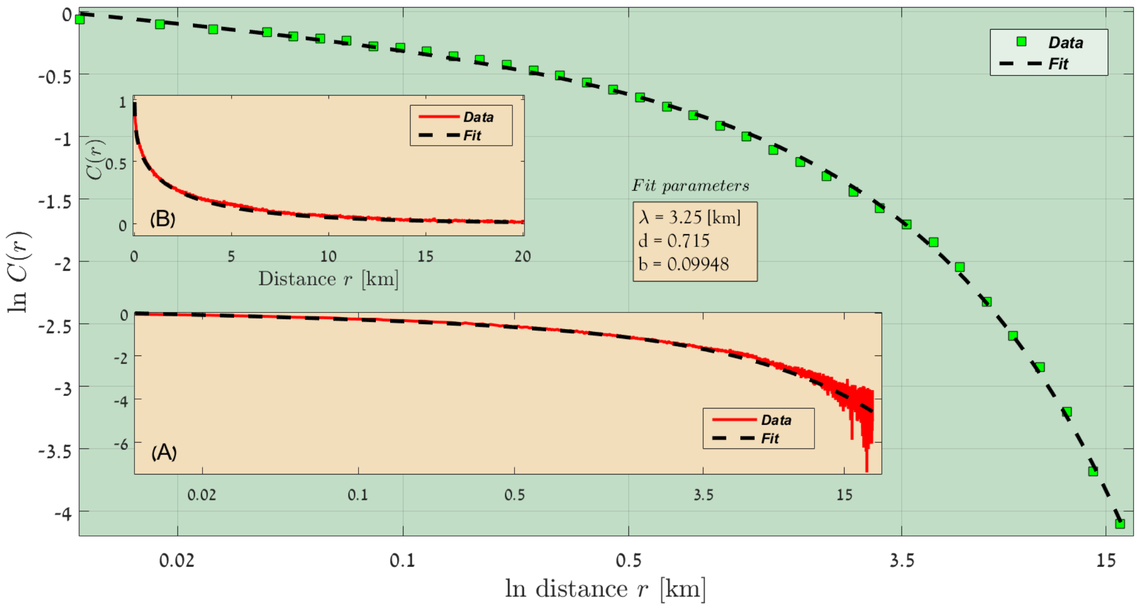


FIG. 17: Sahel, Rainfall lines 850 - 900 [mm/year], 2016-2017.

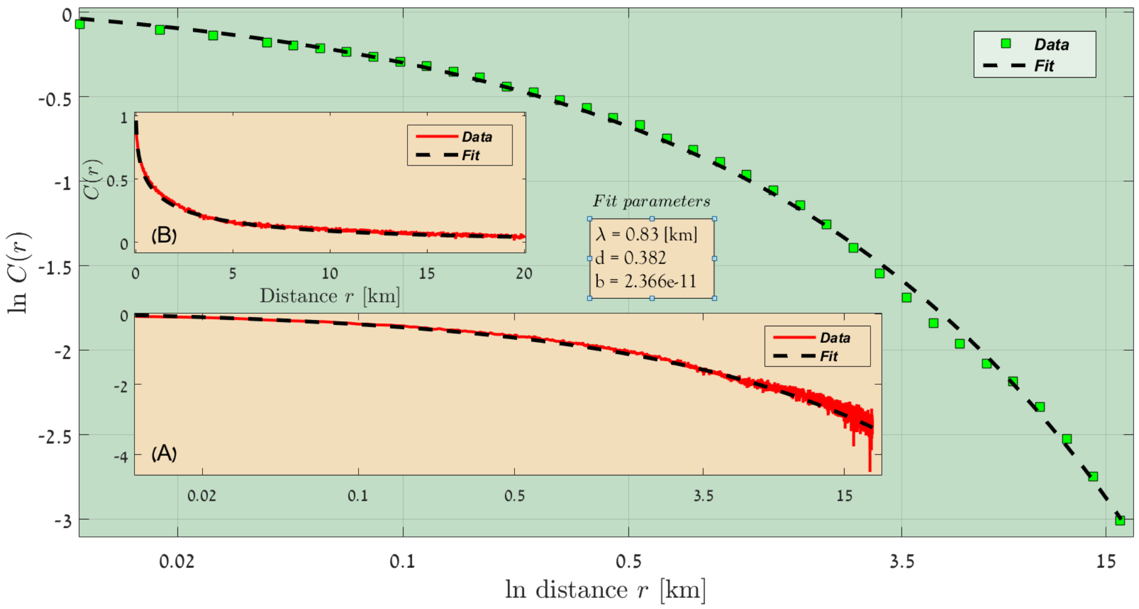


FIG. 18: Sahel, Rainfall lines 900 - 950 [mm/year], 2016-2017.

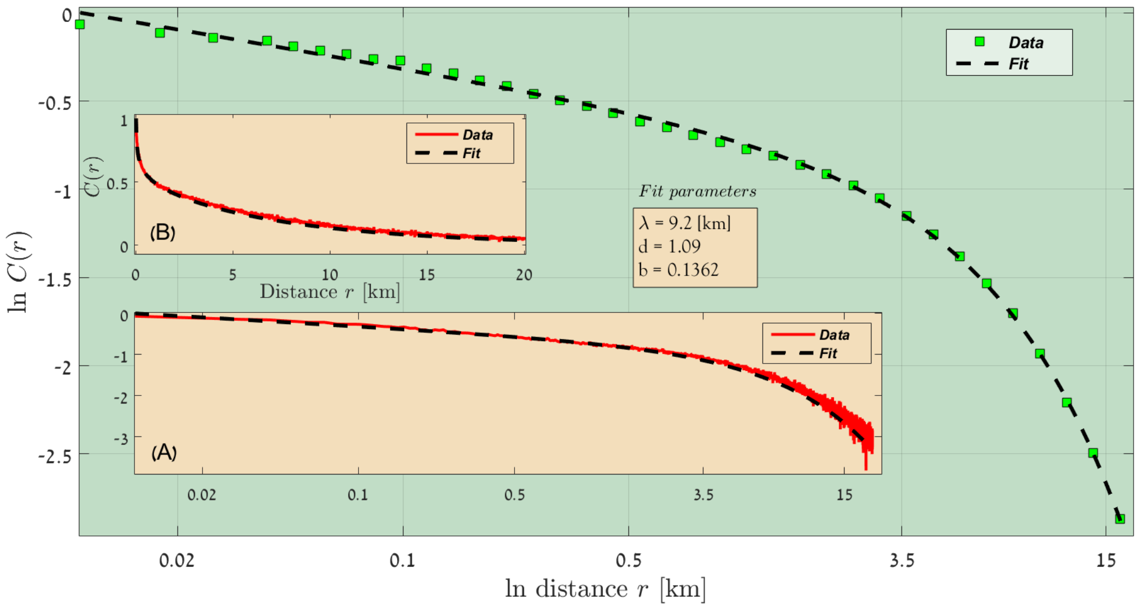


FIG. 19: Sahel, Rainfall lines 950 - 1000 [mm/year], 2016-2017.

### B. Results from Sahel, 13 year lag, 2002-2015

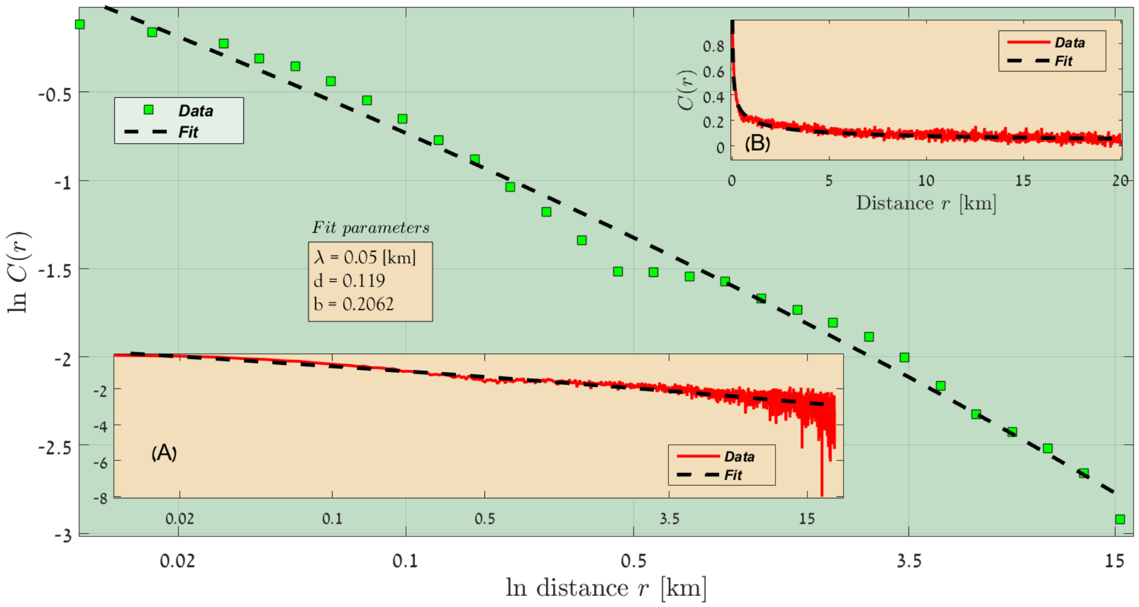


FIG. 20: Sahel, Rainfall lines 200 - 250 [mm/year], 2002-2015.

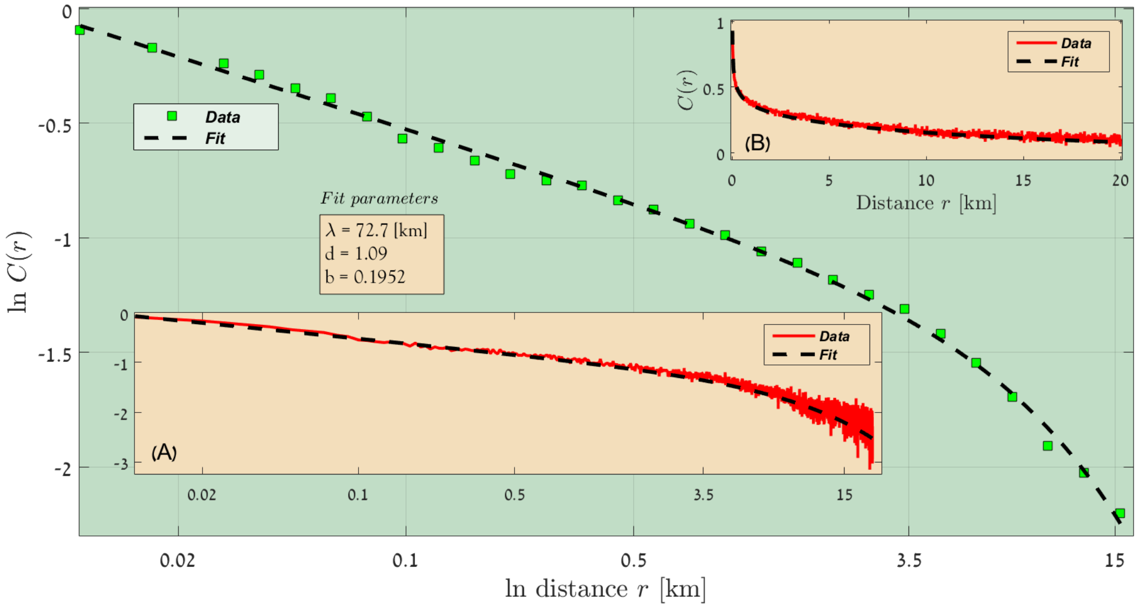


FIG. 21: Sahel, Rainfall lines 250 - 300 [mm/year], 2002-2015.

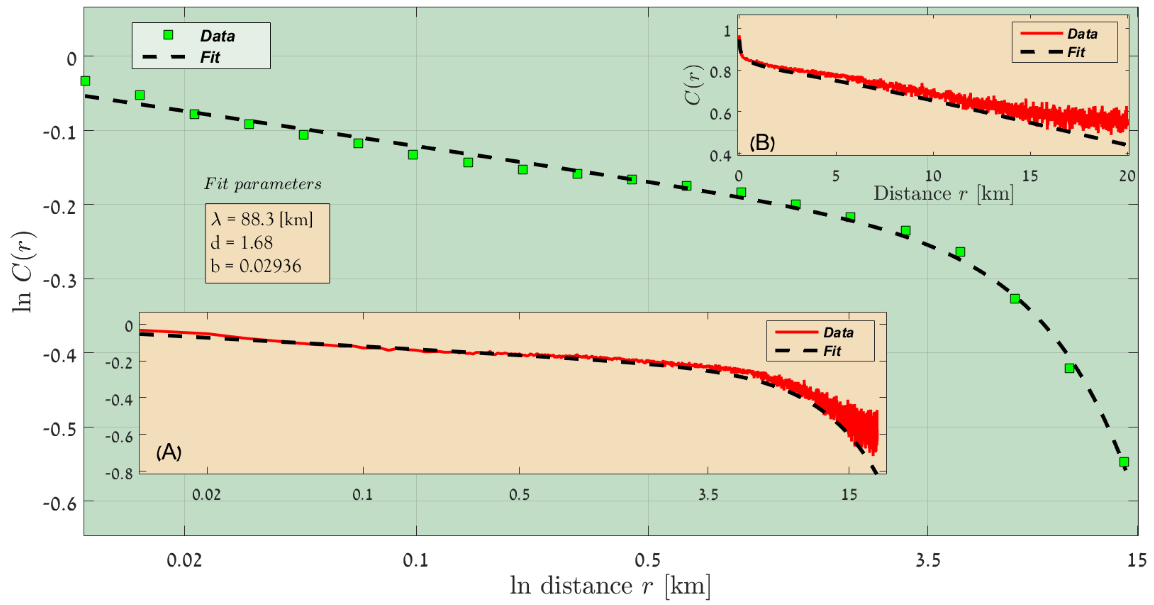


FIG. 22: Sahel, Rainfall lines 300 - 350 [mm/year], 2002-2015.

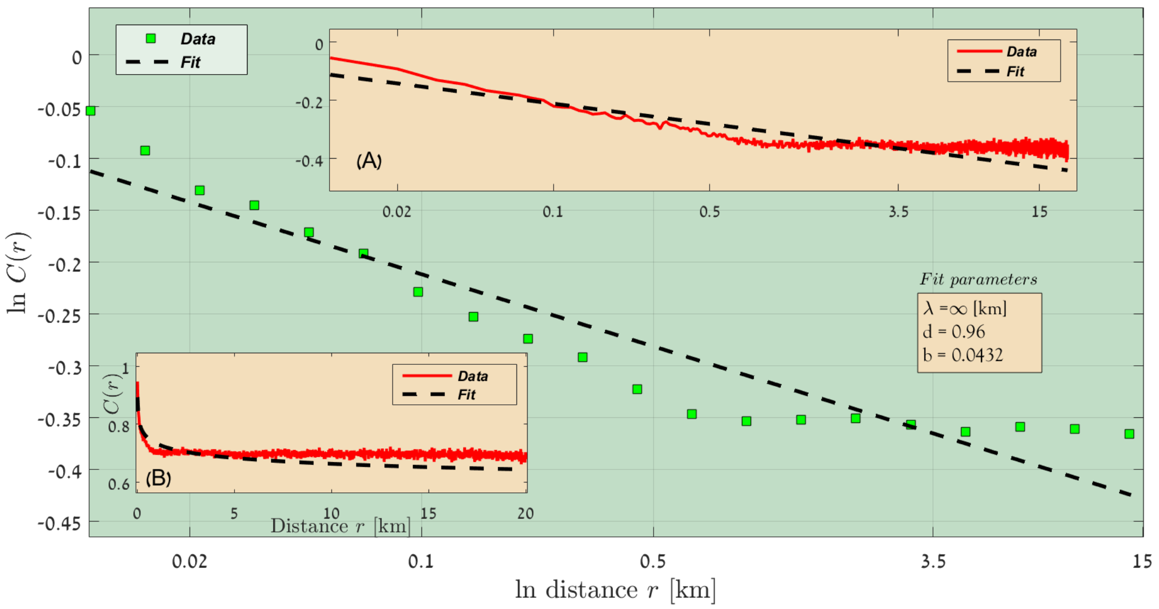


FIG. 23: Sahel, Rainfall lines 350 - 400 [mm/year], 2002-2015.

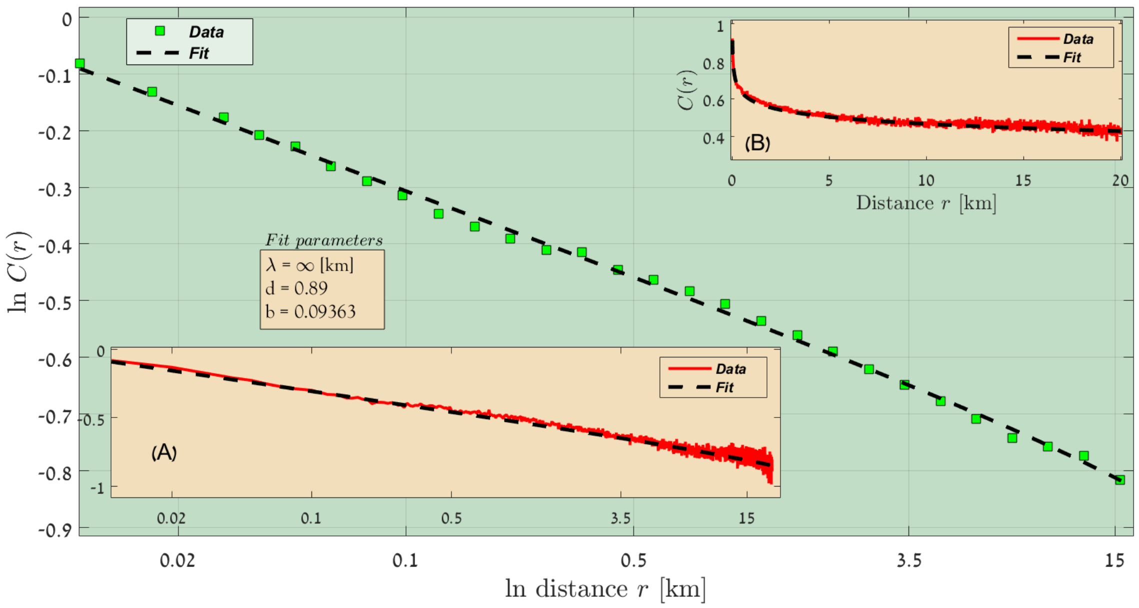


FIG. 24: Sahel, Rainfall lines 400 - 450 [mm/year], 2002-2015.

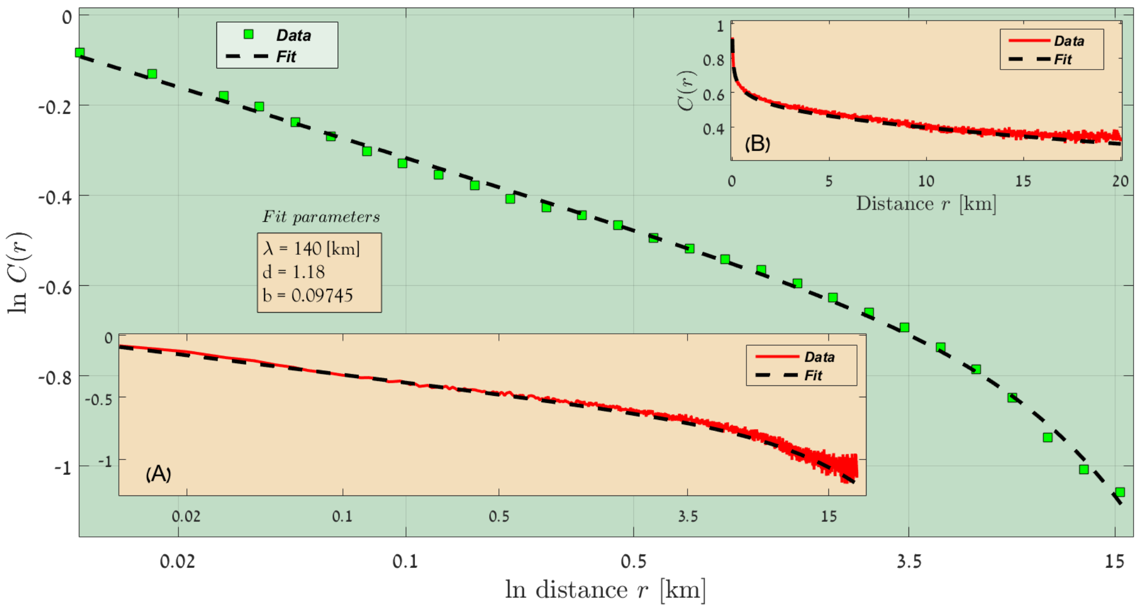


FIG. 25: Sahel, Rainfall lines 450 - 500 [mm/year], 2002-2015.

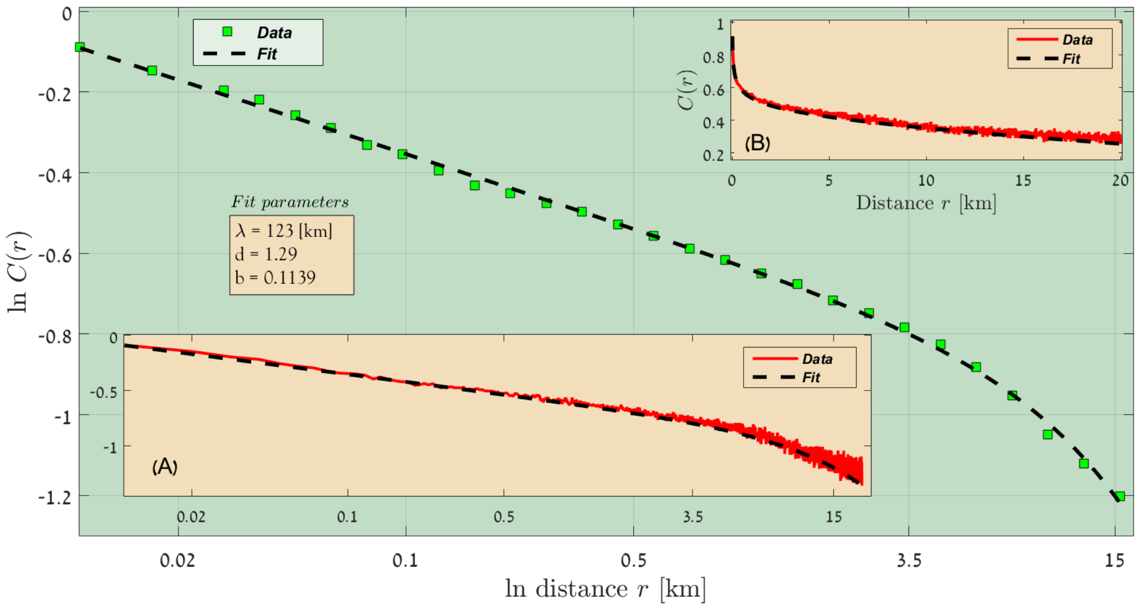


FIG. 26: Sahel, Rainfall lines 500 - 550 [mm/year], 2002-2015.

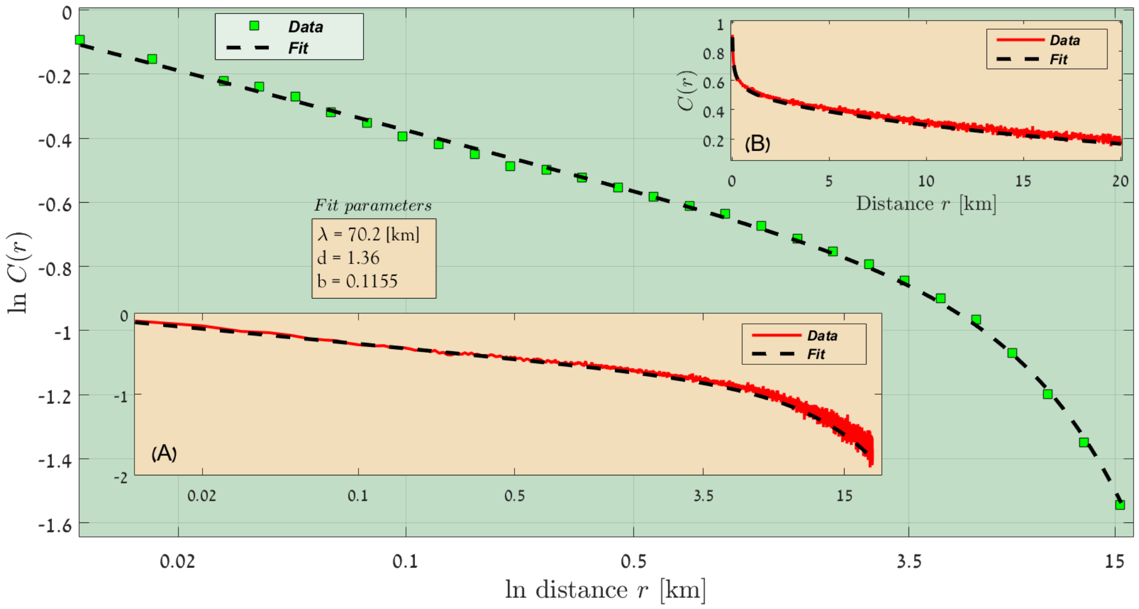


FIG. 27: Sahel, Rainfall lines 5500 - 600 [mm/year], 2002-2015.

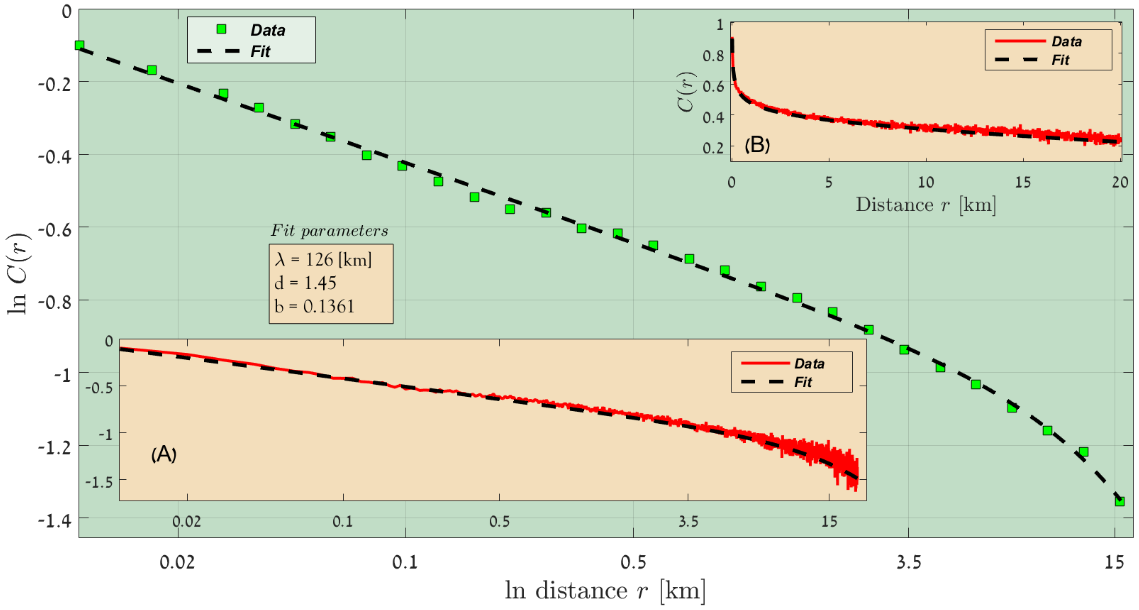


FIG. 28: Sahel, Rainfall lines 600 - 650 [mm/year], 2002-2015.

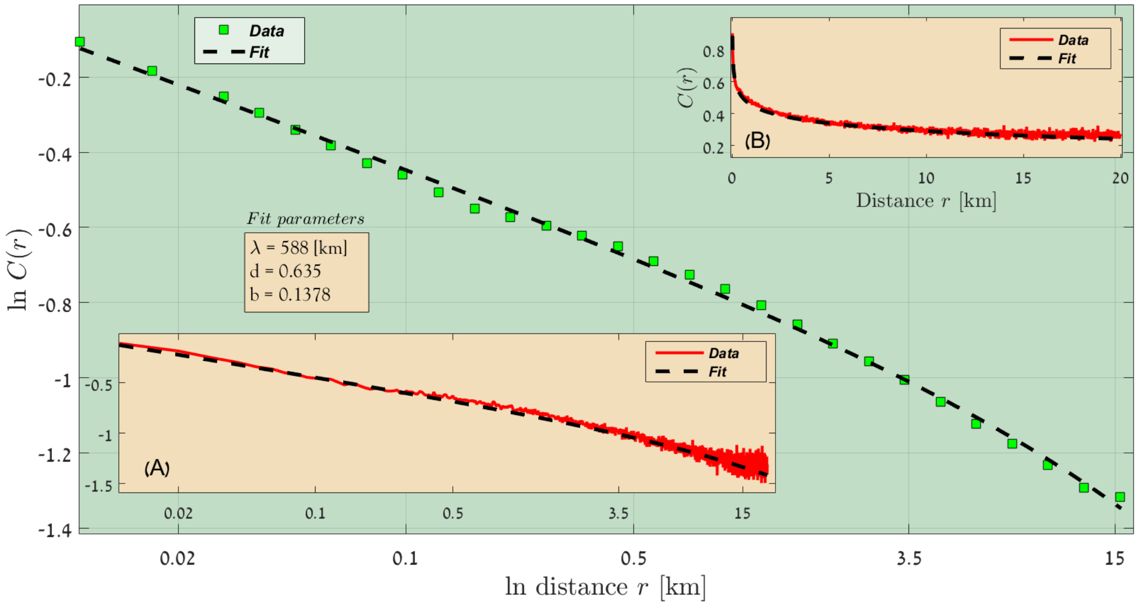


FIG. 29: Sahel, Rainfall lines 650 - 700 [mm/year], 2002-2015.

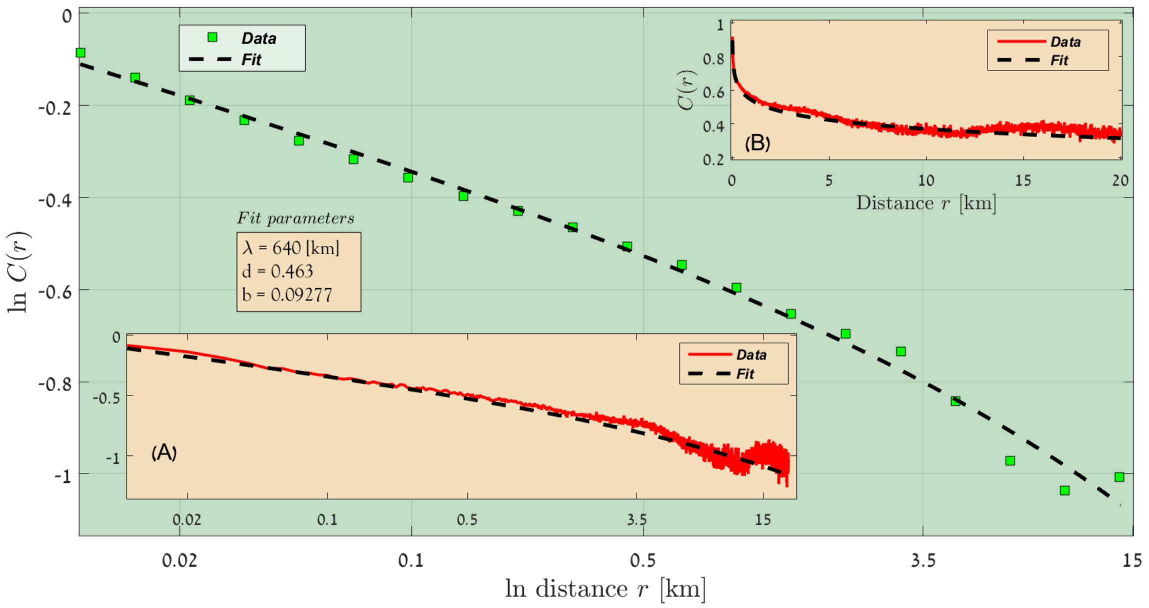


FIG. 30: Sahel, Rainfall lines 700 - 750 [mm/year], 2002-2015.

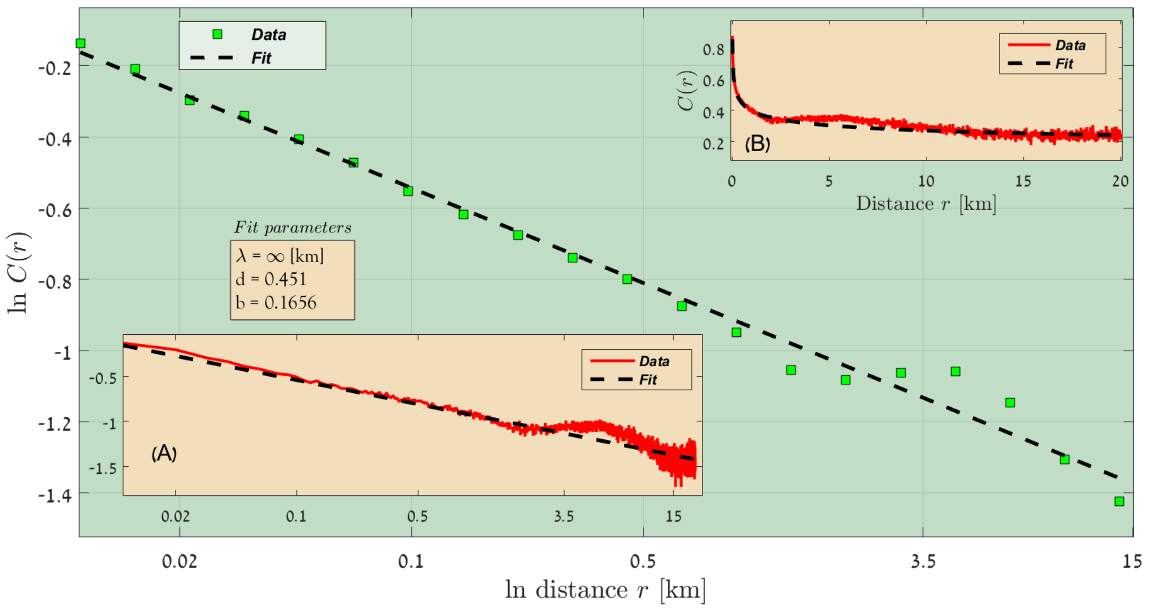


FIG. 31: Sahel, Rainfall lines 750 - 800 [mm/year], 2002-2015.

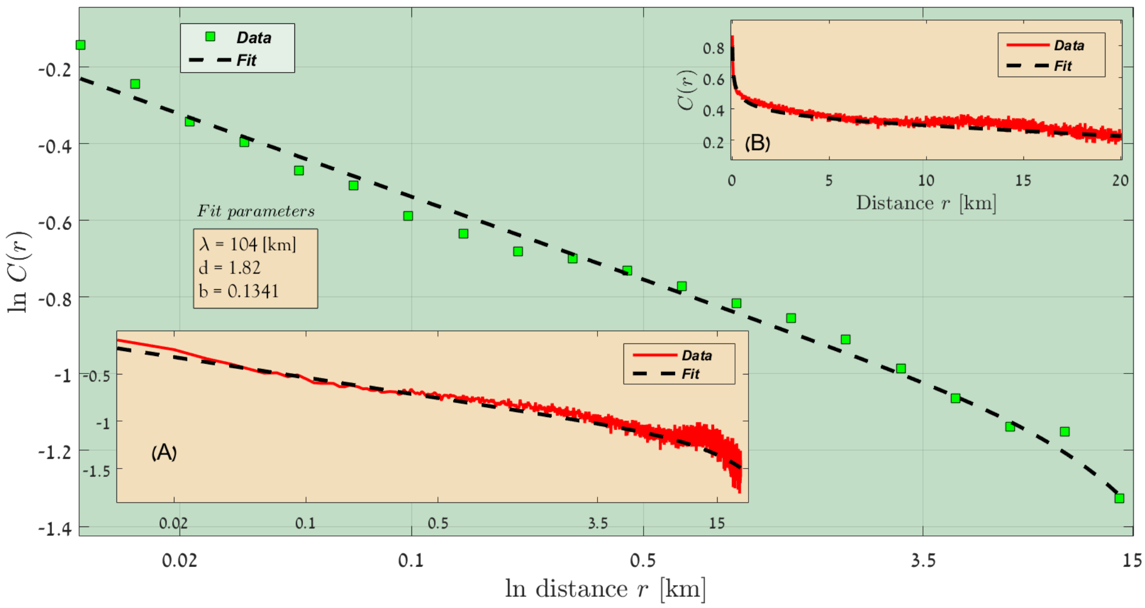


FIG. 32: Sahel, Rainfall lines 750 - 800 [mm/year], 2002-2015.

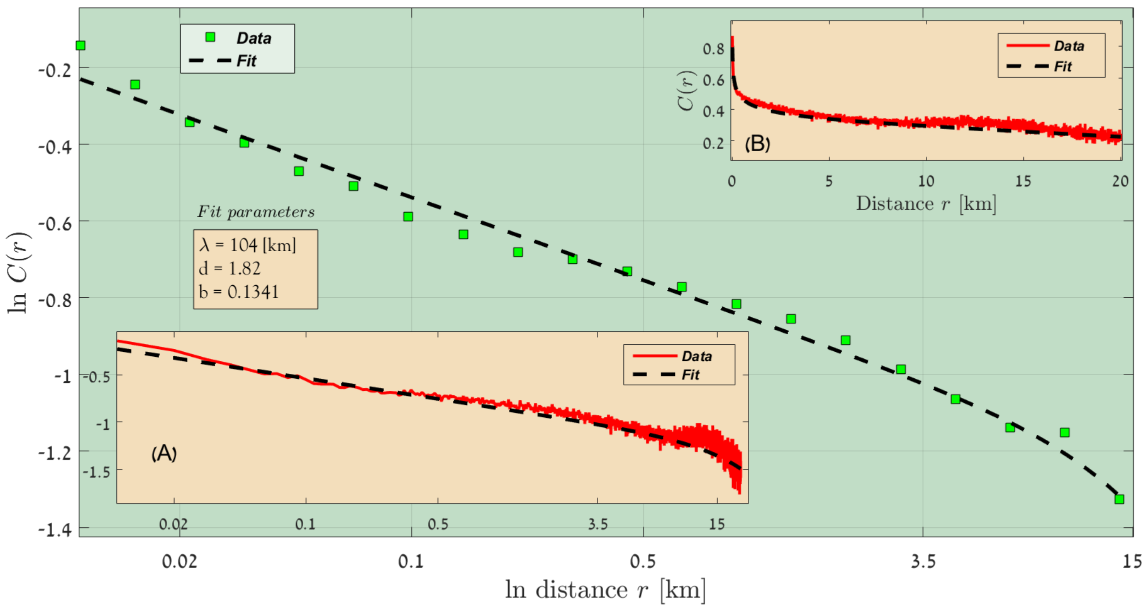


FIG. 33: Sahel, Rainfall lines 800 - 850 [mm/year], 2002-2015.

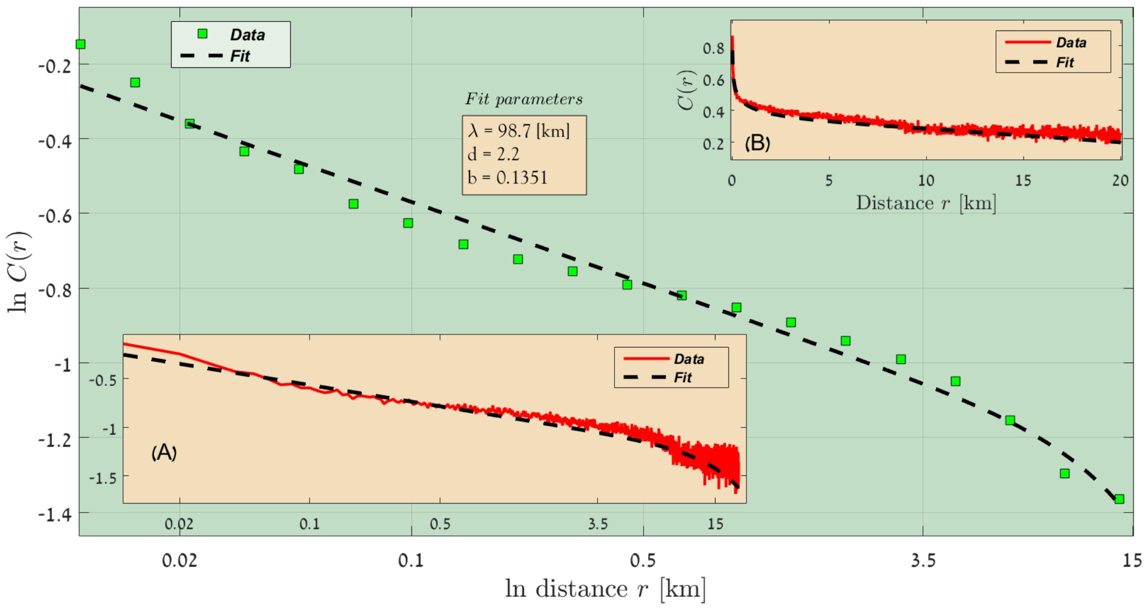


FIG. 34: Sahel, Rainfall lines 850 - 900 [mm/year], 2002-2015.

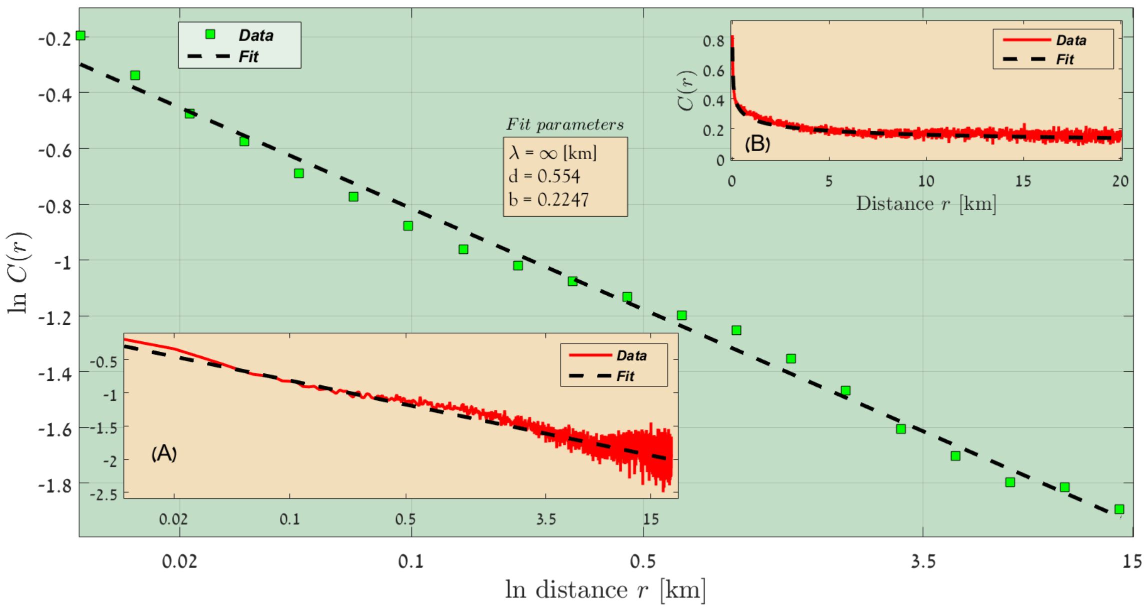


FIG. 35: Sahel, Rainfall lines 900 - 950 [mm/year], 2002-2015.

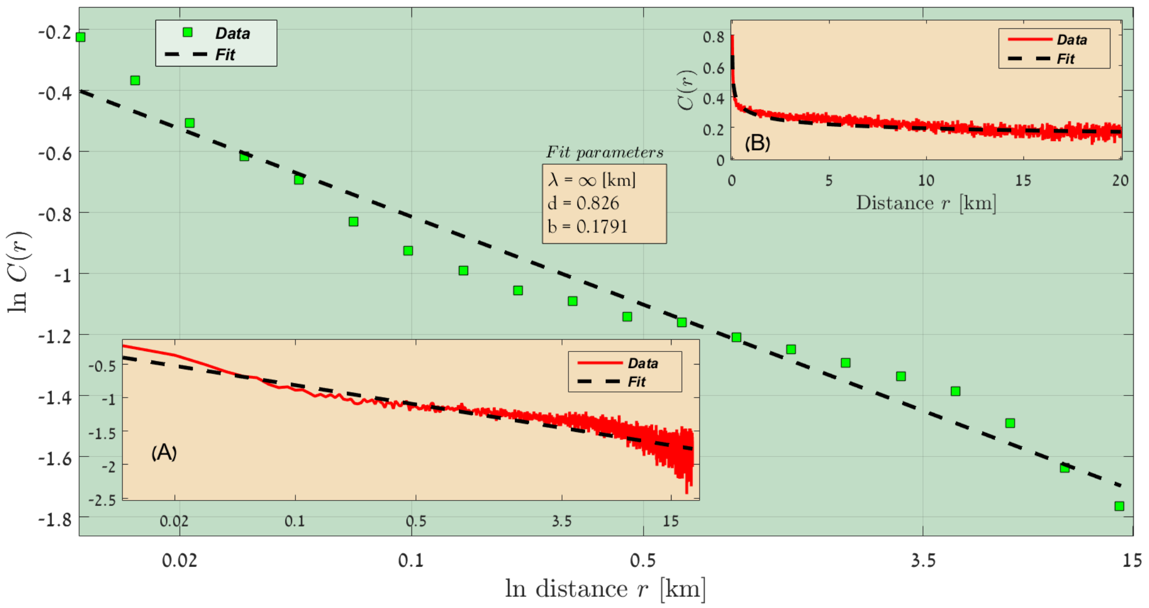


FIG. 36: Sahel, Rainfall lines 950 - 1000 [mm/year], 2002-2015.

#### C. Results from Australia, 1-2 year lag

**Australia, 2016-2017**

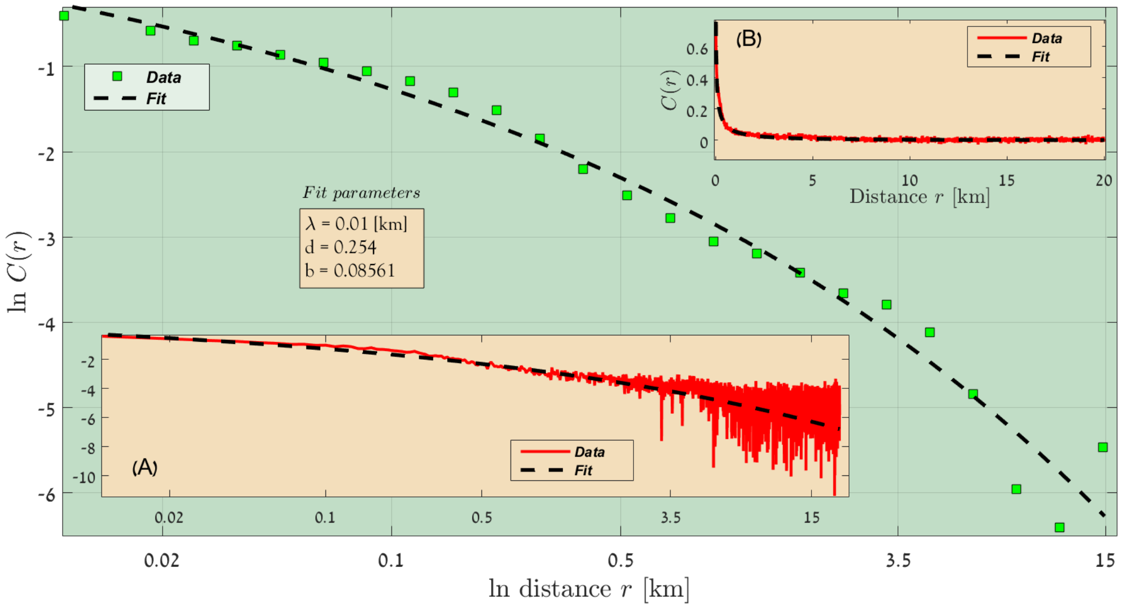


FIG. 37: Australia, Rainfall lines 50 - 100 [mm/year], 2016-2017.

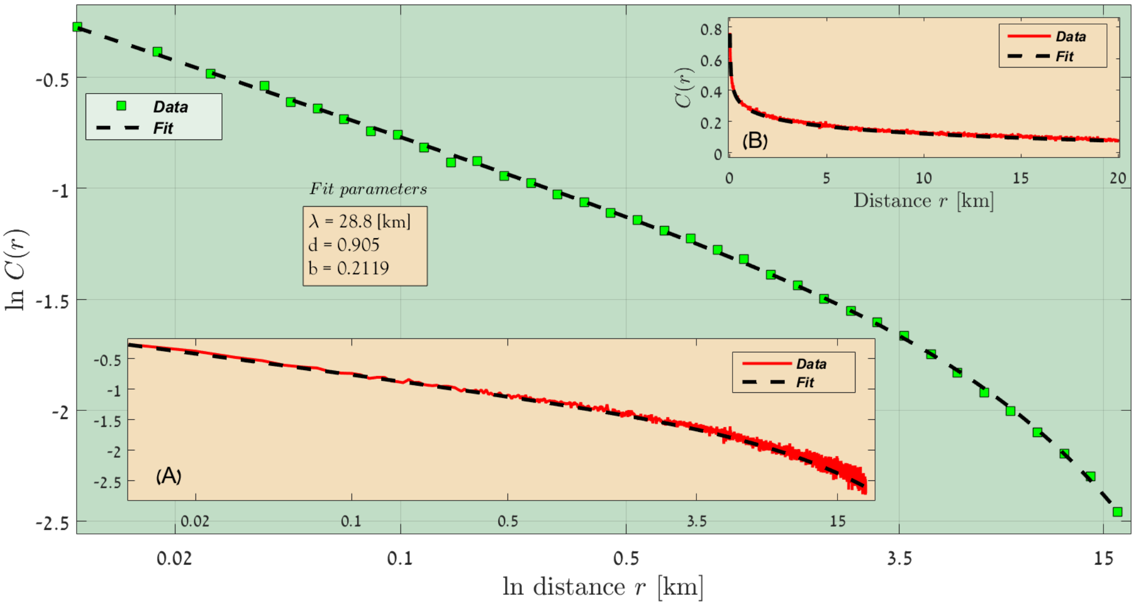


FIG. 38: Australia, Rainfall lines 100 - 150 [mm/year], 2016-2017.

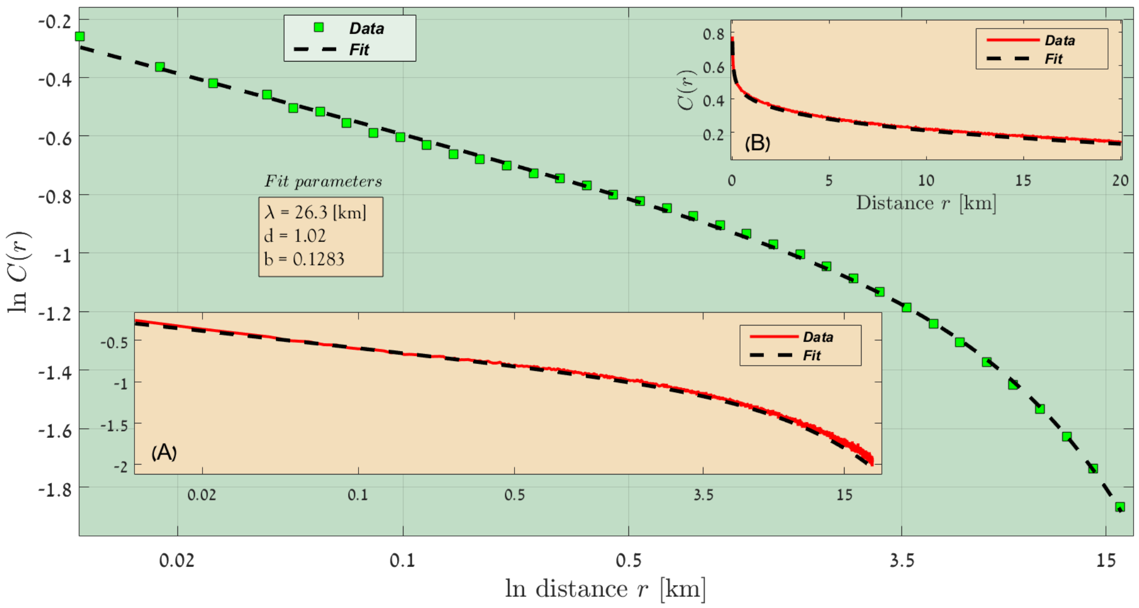


FIG. 39: Australia, Rainfall lines 150 - 200 [mm/year], 2016-2017.

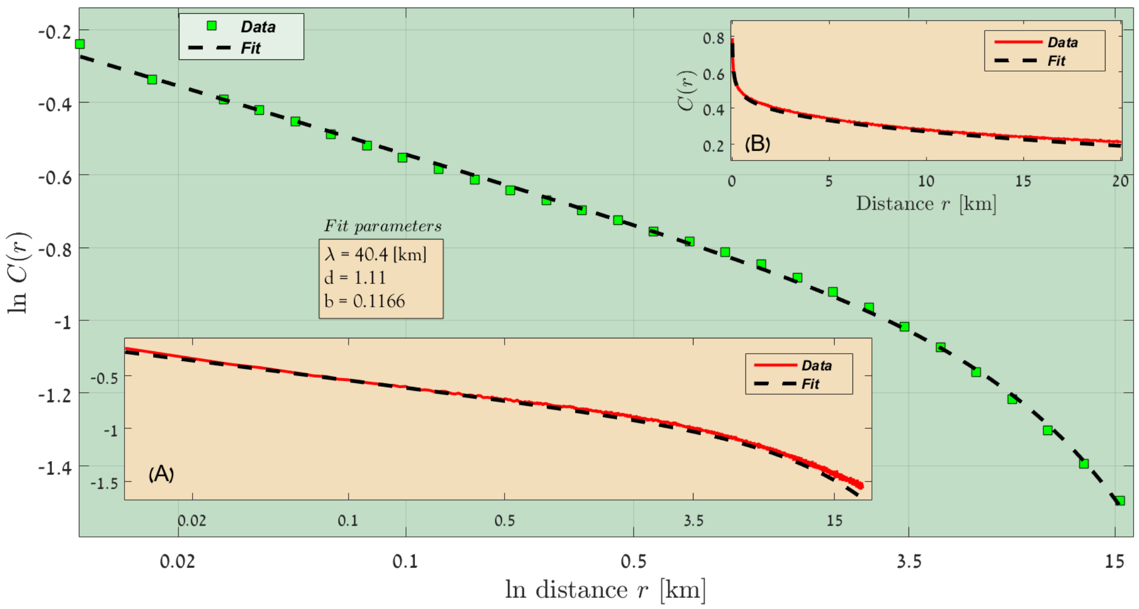


FIG. 40: Australia, Rainfall lines 200 - 250 [mm/year], 2016-2017.

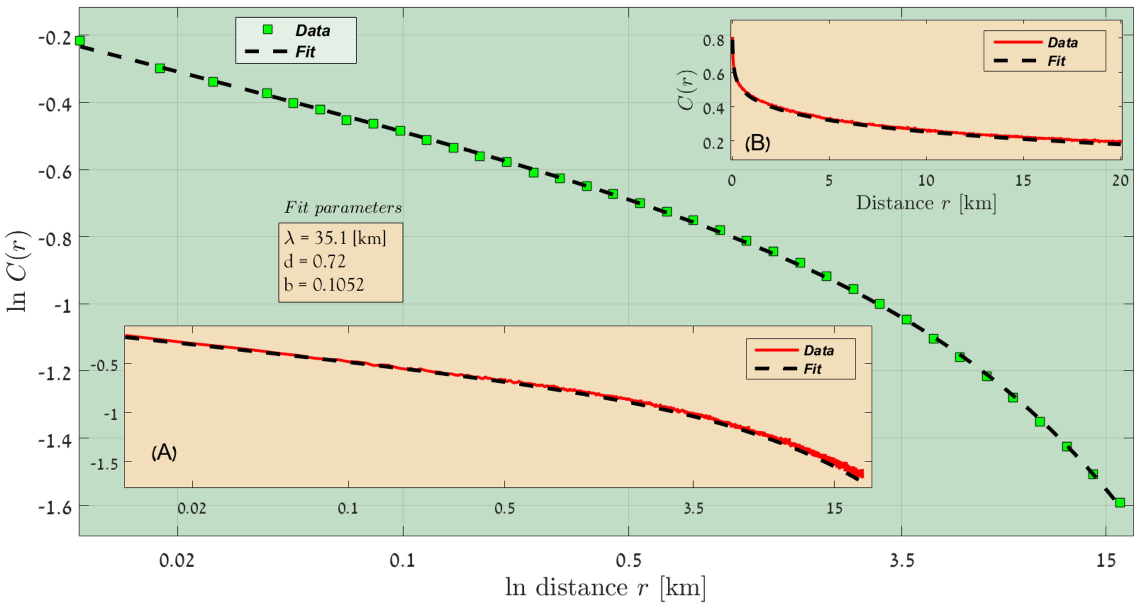


FIG. 41: Australia, Rainfall lines 250 - 300 [mm/year], 2016-2017.

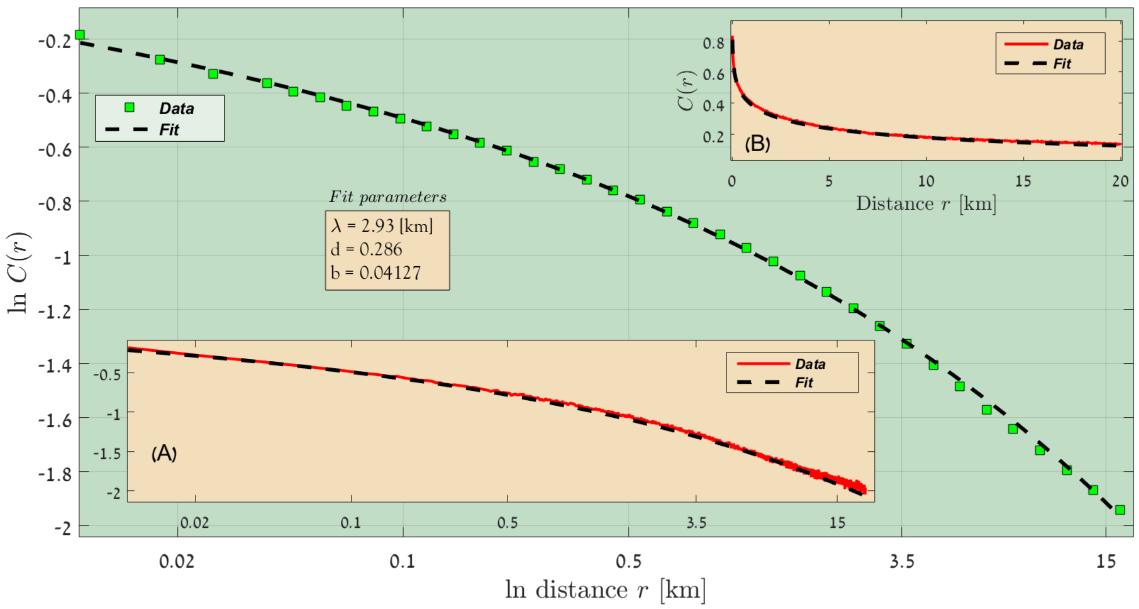


FIG. 42: Australia, Rainfall lines 300 - 350 [mm/year], 2016-2017.

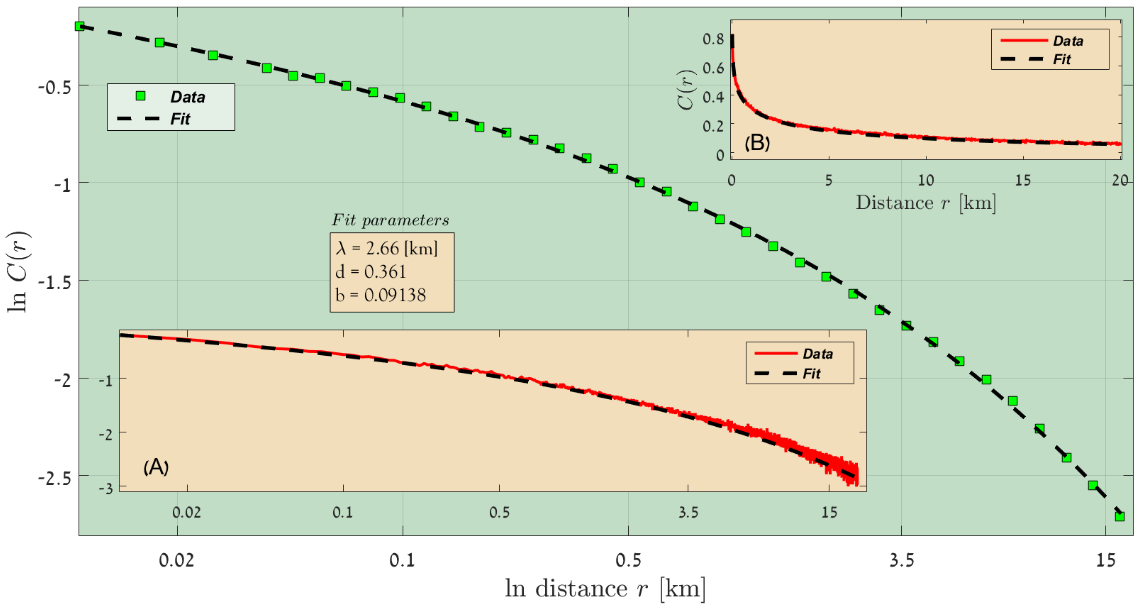


FIG. 43: Australia, Rainfall lines 350 - 400 [mm/year], 2016-2017.

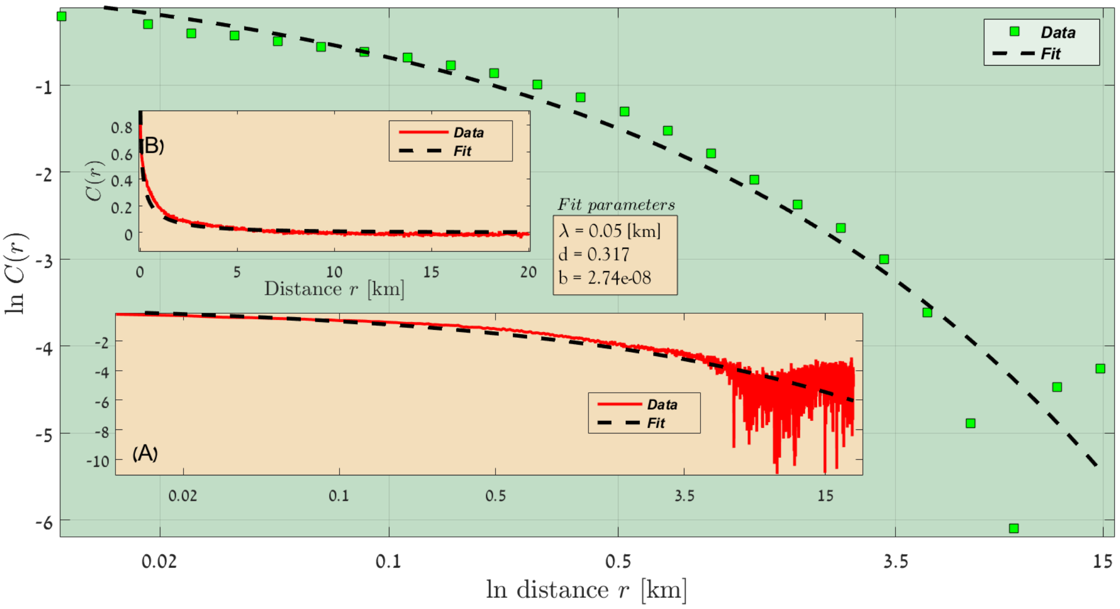


FIG. 44: Australia, Rainfall lines 400 - 450 [mm/year], 2016-2017.

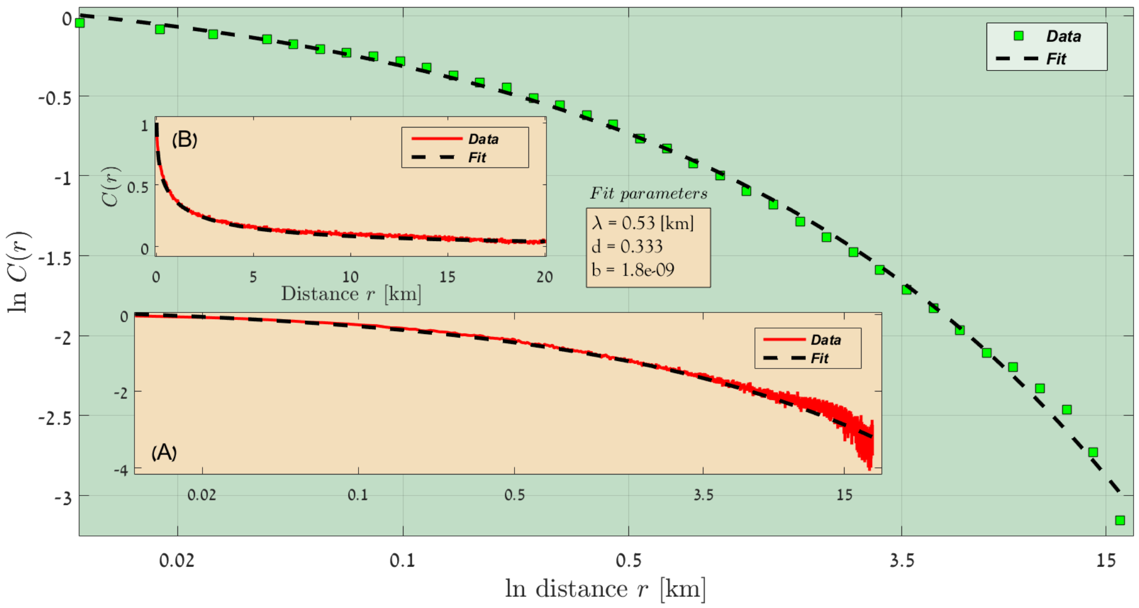


FIG. 45: Australia, Rainfall lines 450 - 500 [mm/year], 2016-2017.

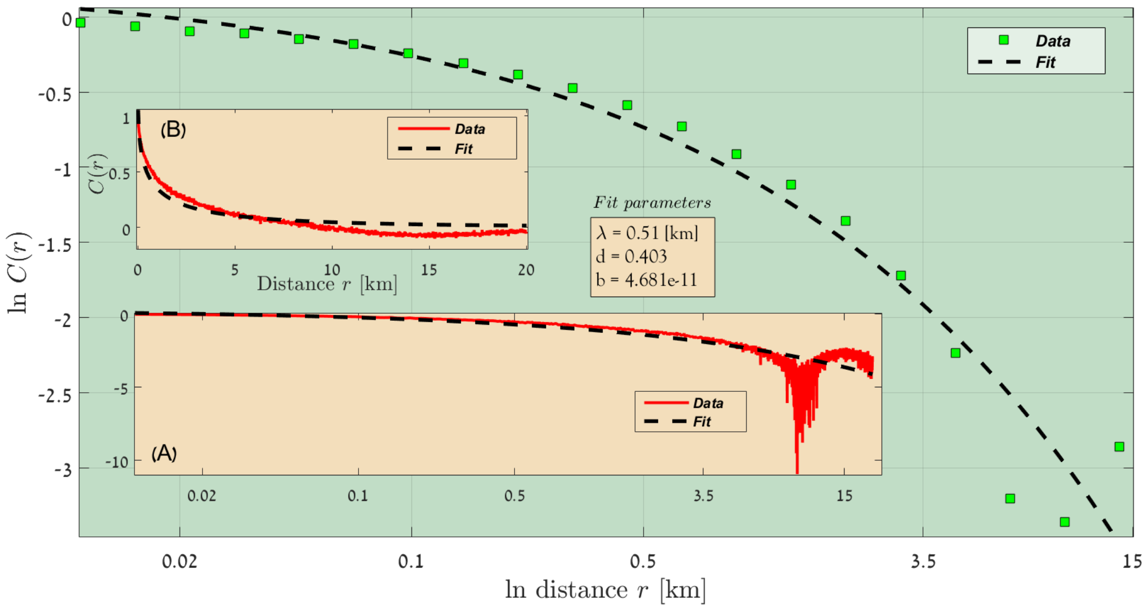


FIG. 46: Australia, Rainfall lines 500 - 550 [mm/year], 2016-2017.

**Australia, 2017-2018**

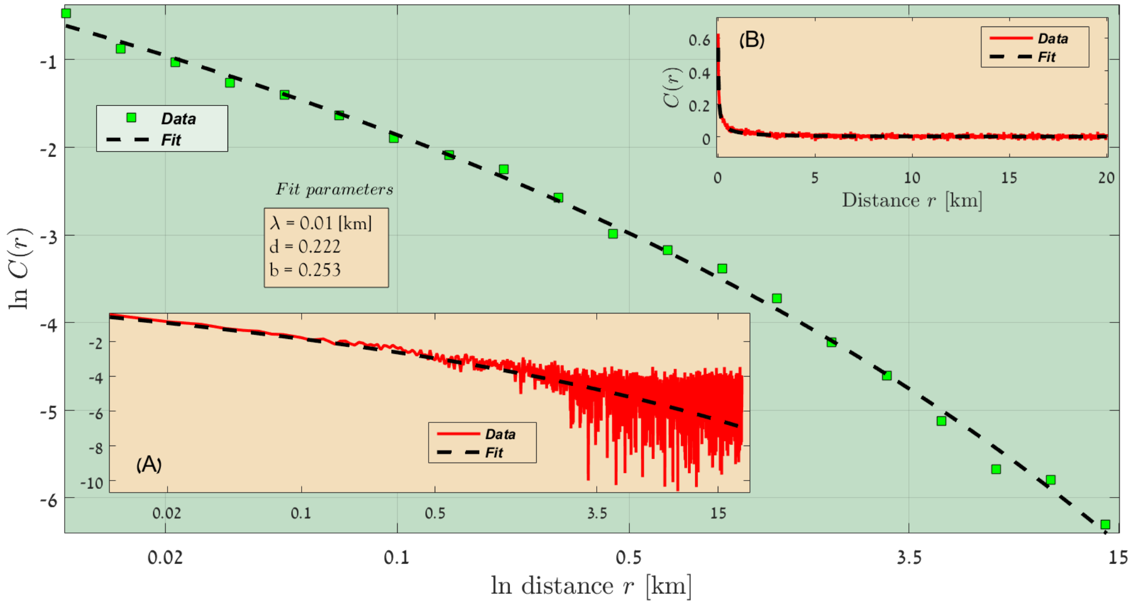


FIG. 47: Australia, Rainfall lines 50 - 100 [mm/year], 2017-2018.

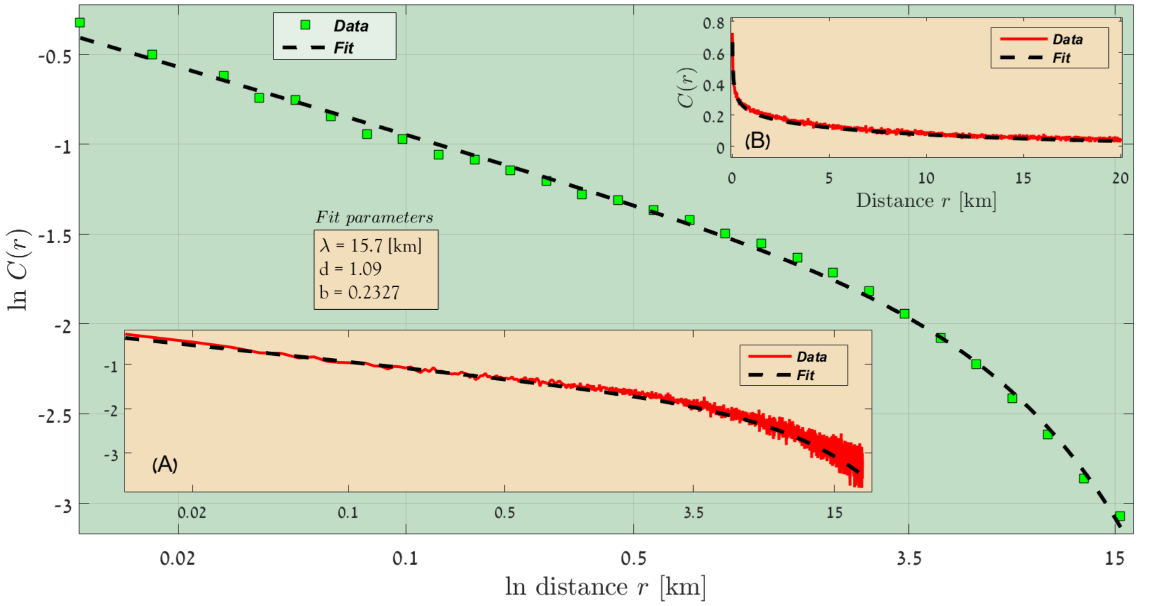


FIG. 48: Australia, Rainfall lines 100 - 150 [mm/year], 2017-2018.

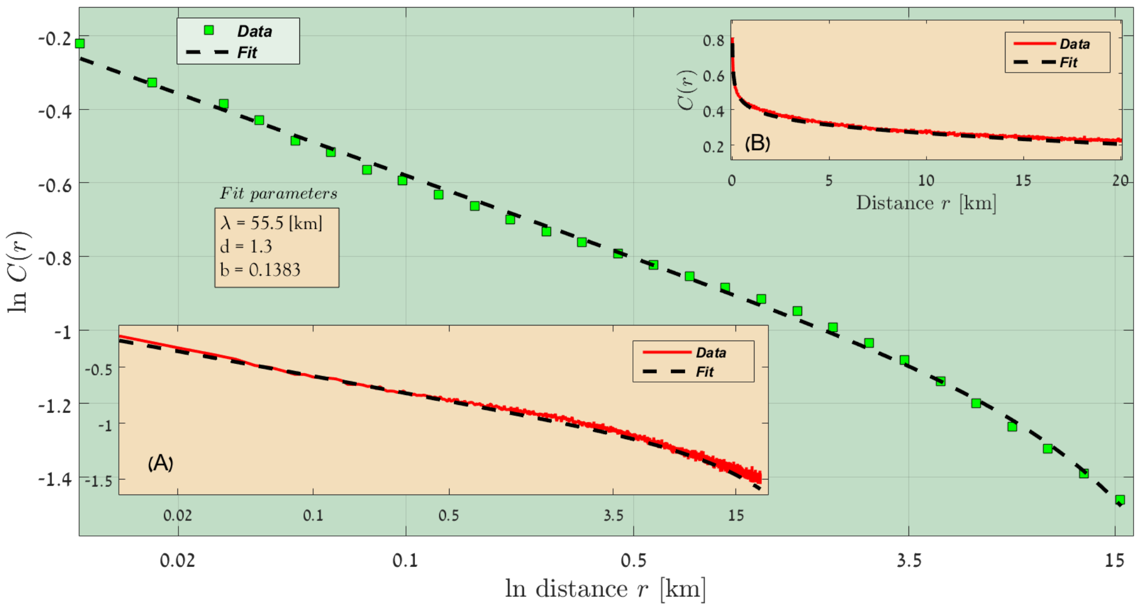


FIG. 49: Australia, Rainfall lines 150 - 200 [mm/year], 2017-2018.

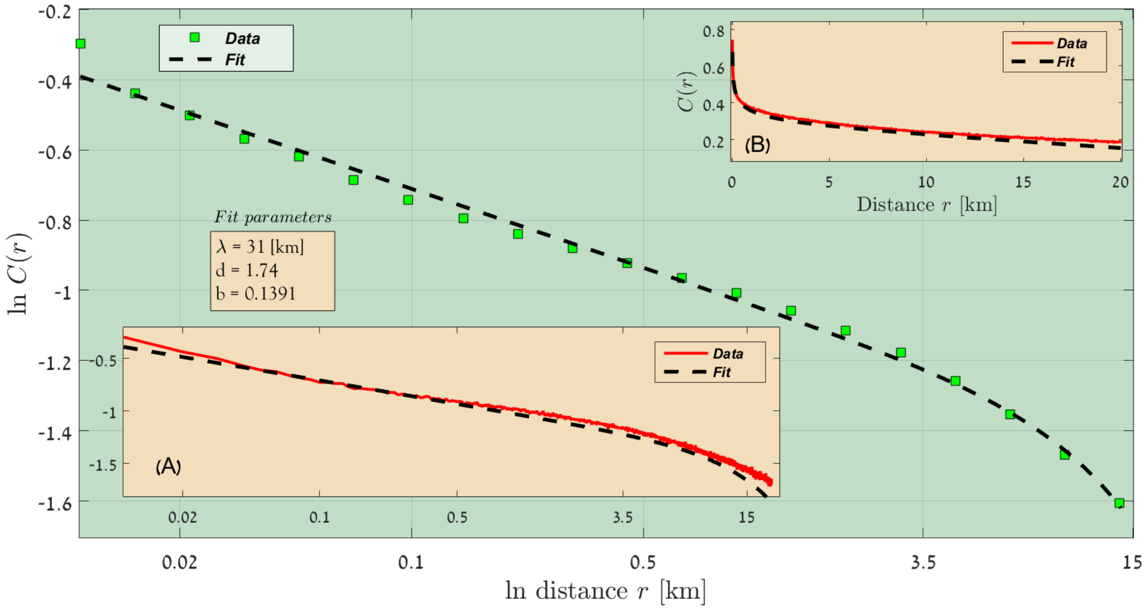


FIG. 50: Australia, Rainfall lines 200 - 250 [mm/year], 2017-2018.

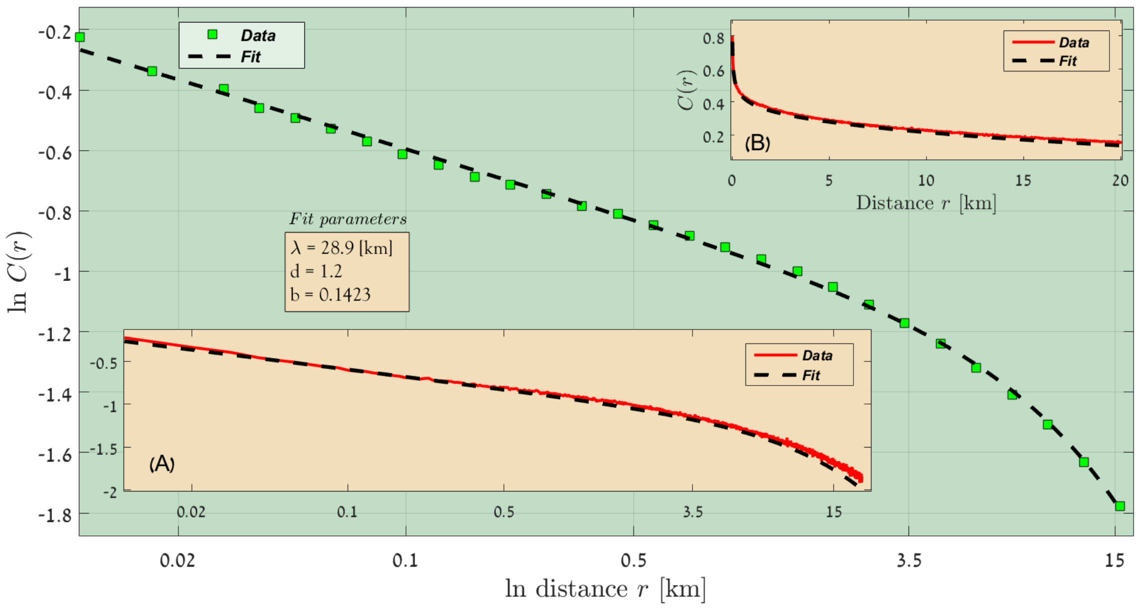


FIG. 51: Australia, Rainfall lines 250 - 300 [mm/year], 2017-2018.

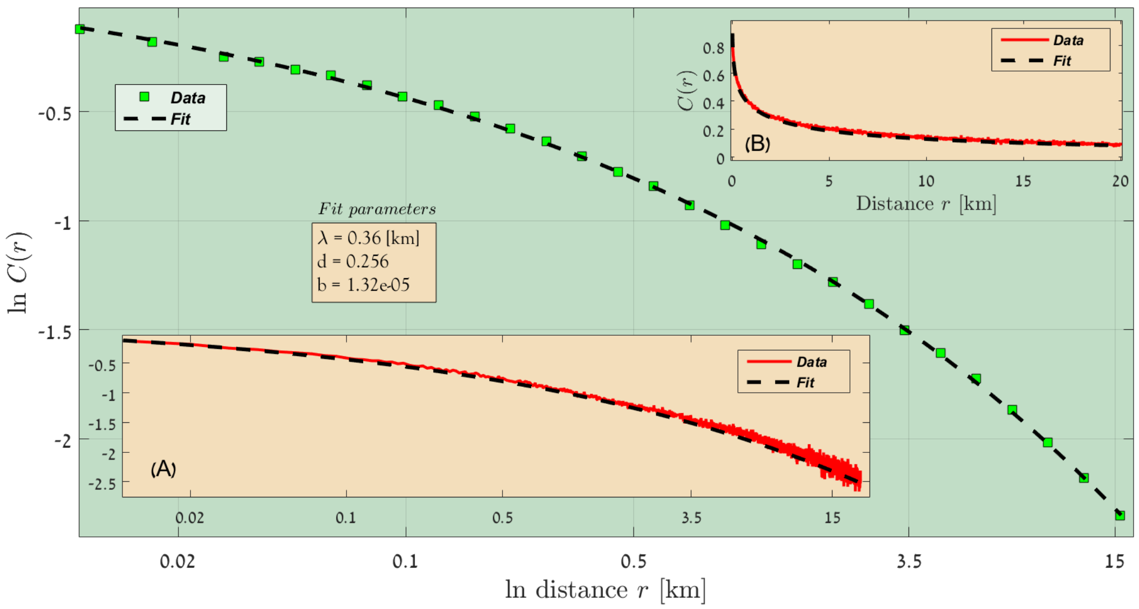


FIG. 52: Australia, Rainfall lines 300 - 350 [mm/year], 2017-2018.

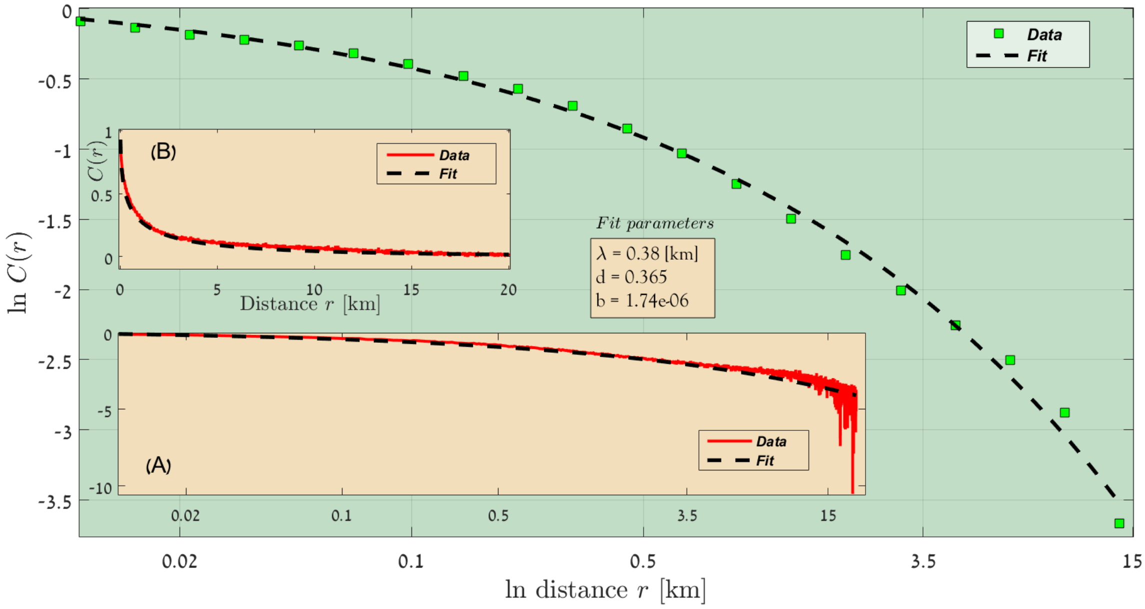


FIG. 53: Australia, Rainfall lines 350 - 400 [mm/year], 2017-2018.

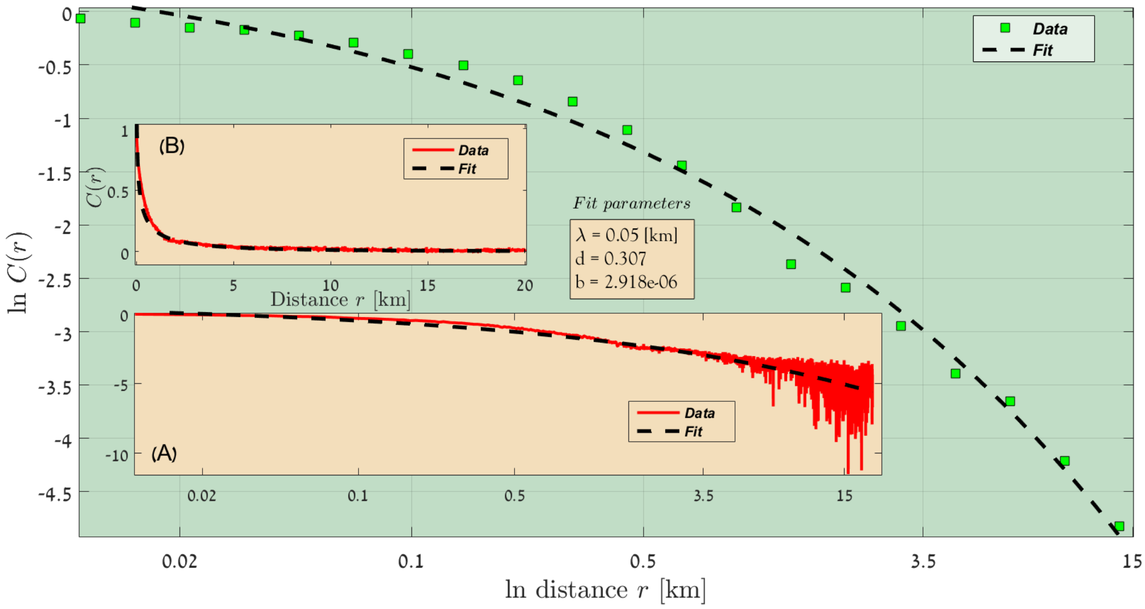


FIG. 54: Australia, Rainfall lines 400 - 450 [mm/year], 2017-2018.

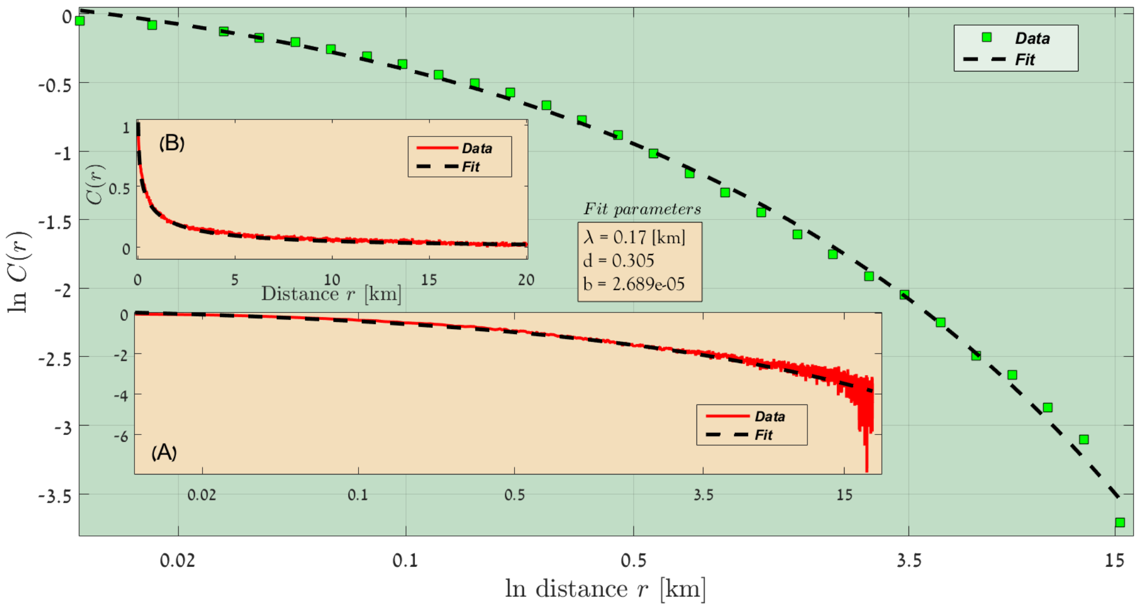


FIG. 55: Australia, Rainfall lines 450 - 500 [mm/year], 2017-2018.

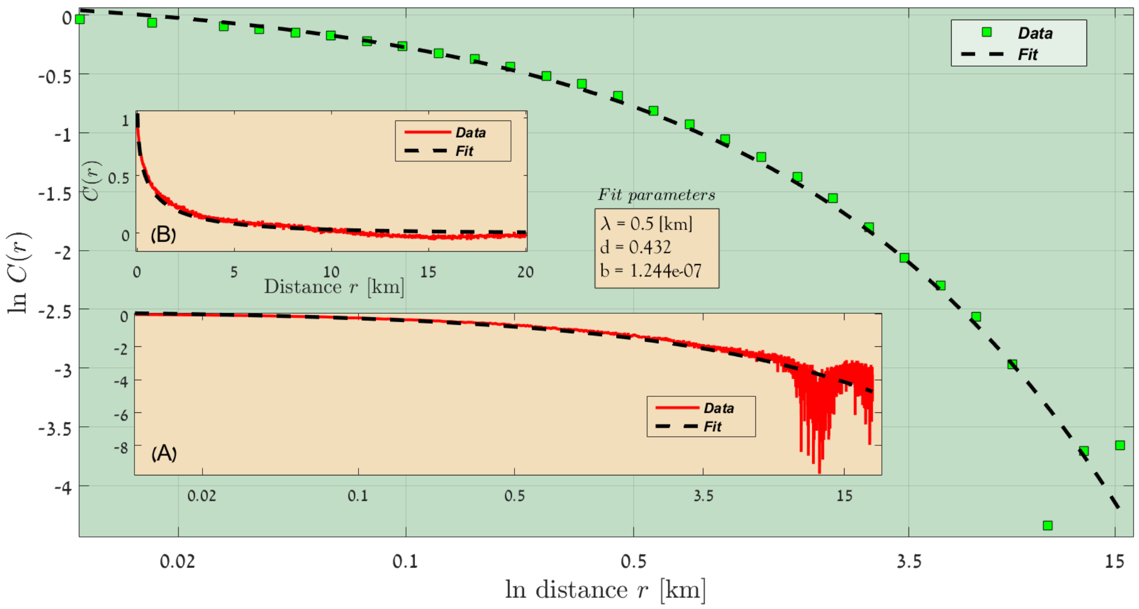


FIG. 56: Australia, Rainfall lines 500 - 550 [mm/year], 2017-2018.

**Australia, 2016-2018**

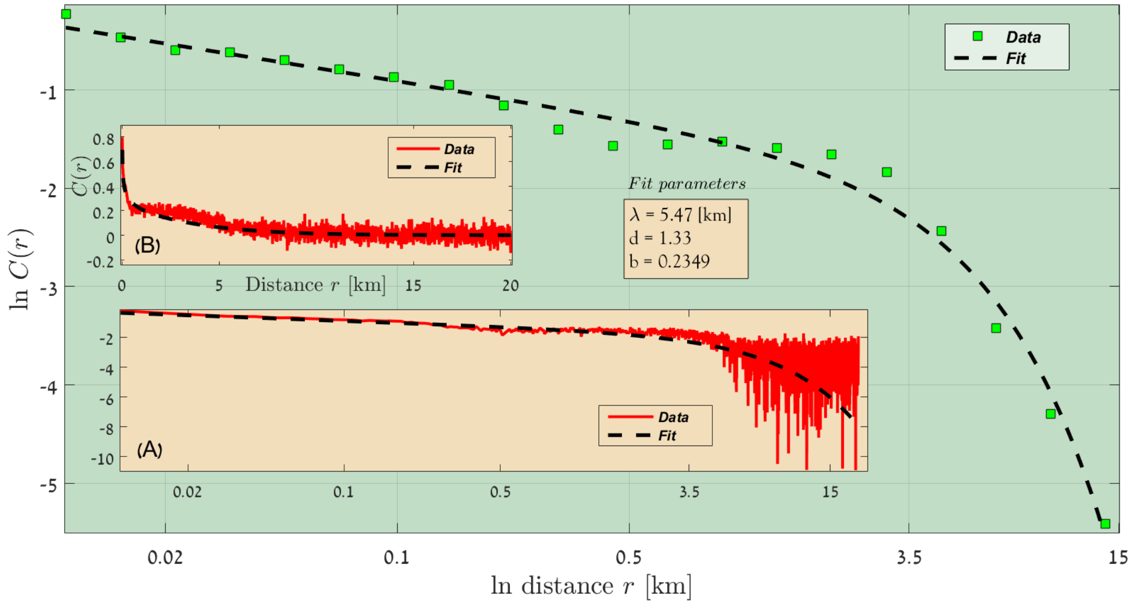


FIG. 57: Australia, Rainfall lines 50 - 100 [mm/year], 2016-2018.

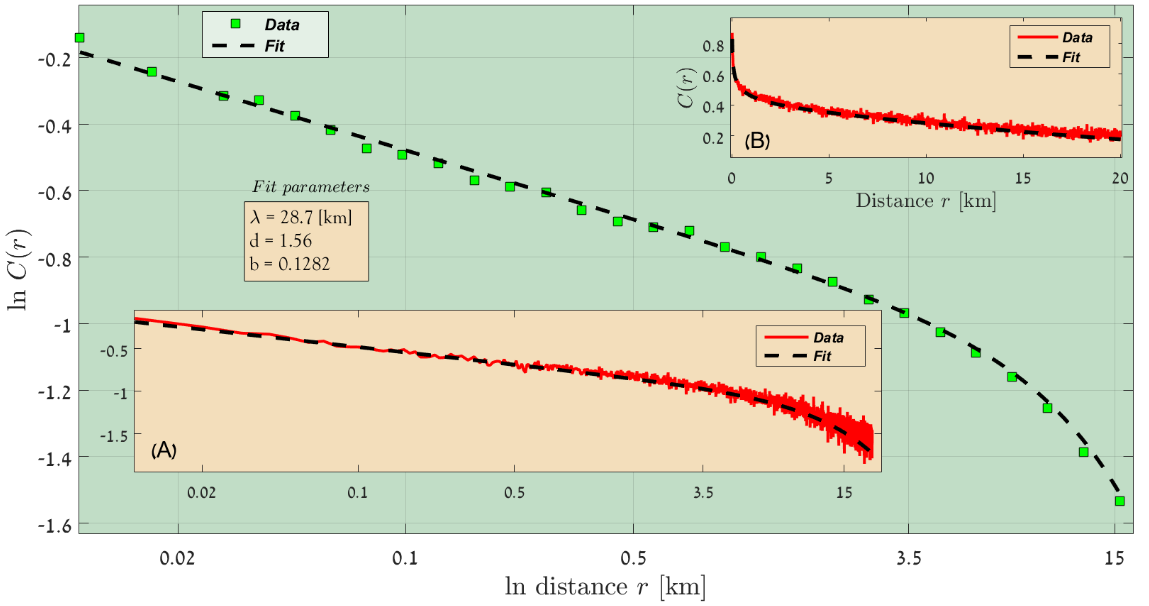


FIG. 58: Australia, Rainfall lines 100 - 150 [mm/year], 2016-2018.

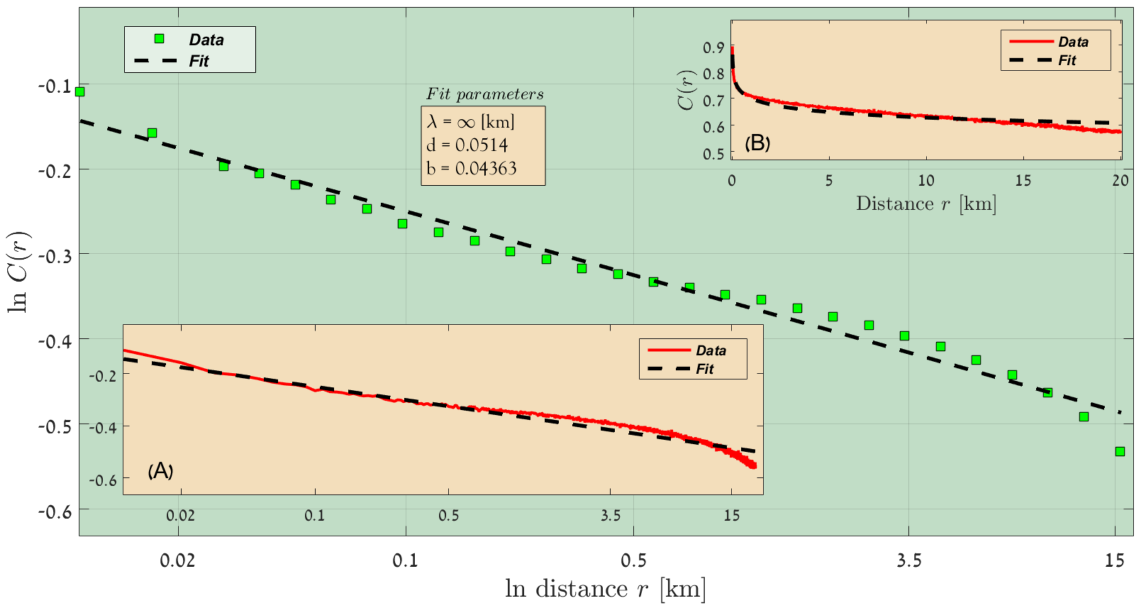


FIG. 59: Australia, Rainfall lines 150 - 200 [mm/year], 2016-2018.

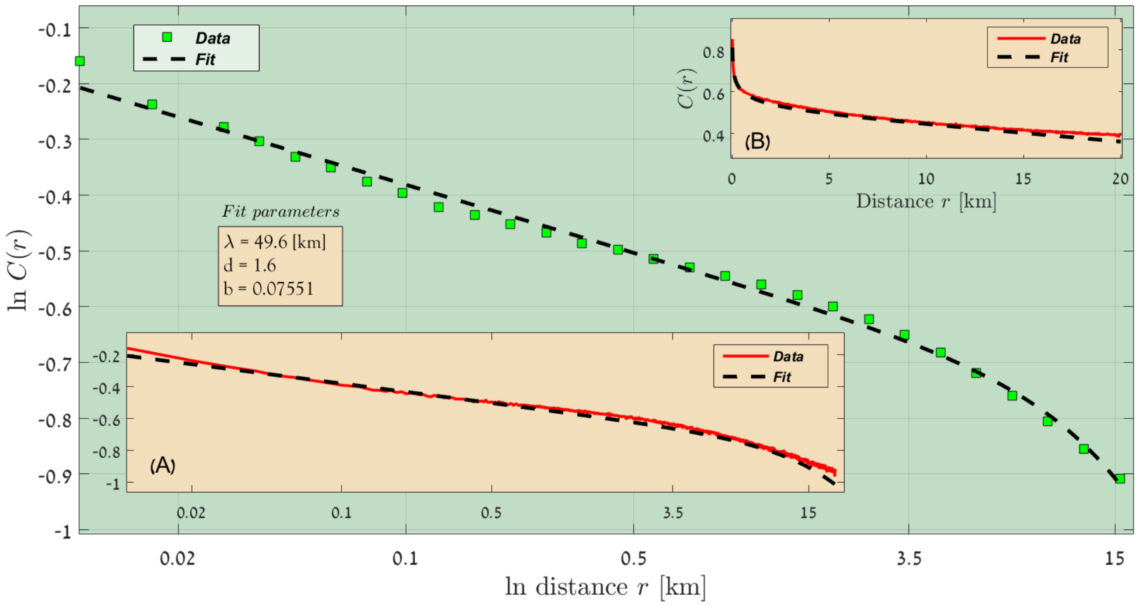


FIG. 60: Australia, Rainfall lines 200 - 250 [mm/year], 2016-2018.

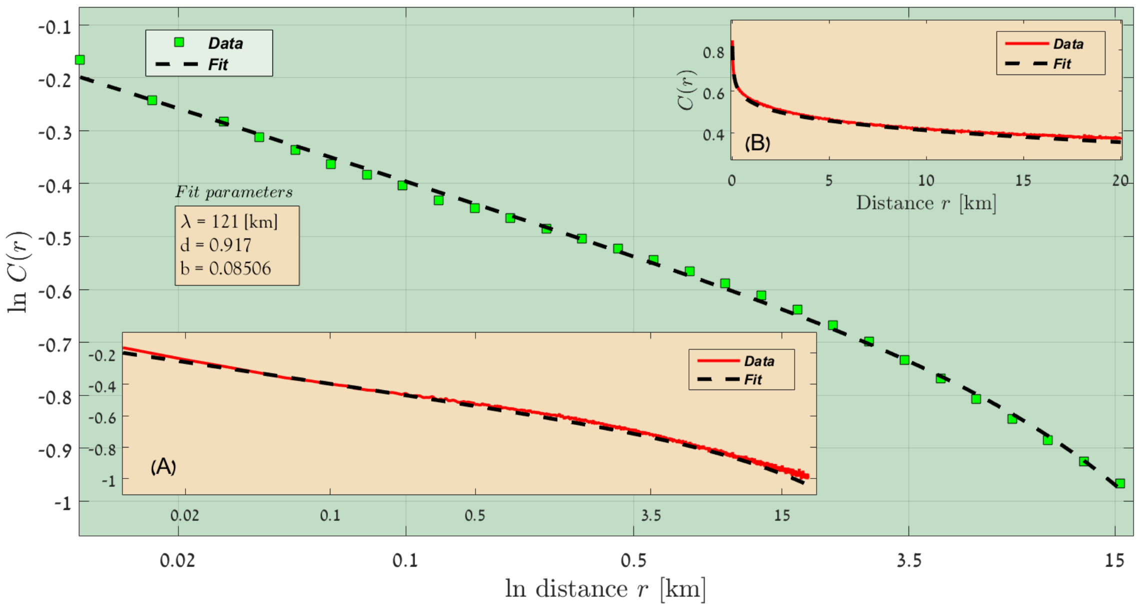


FIG. 61: Australia, Rainfall lines 250 - 300 [mm/year], 2016-2018.

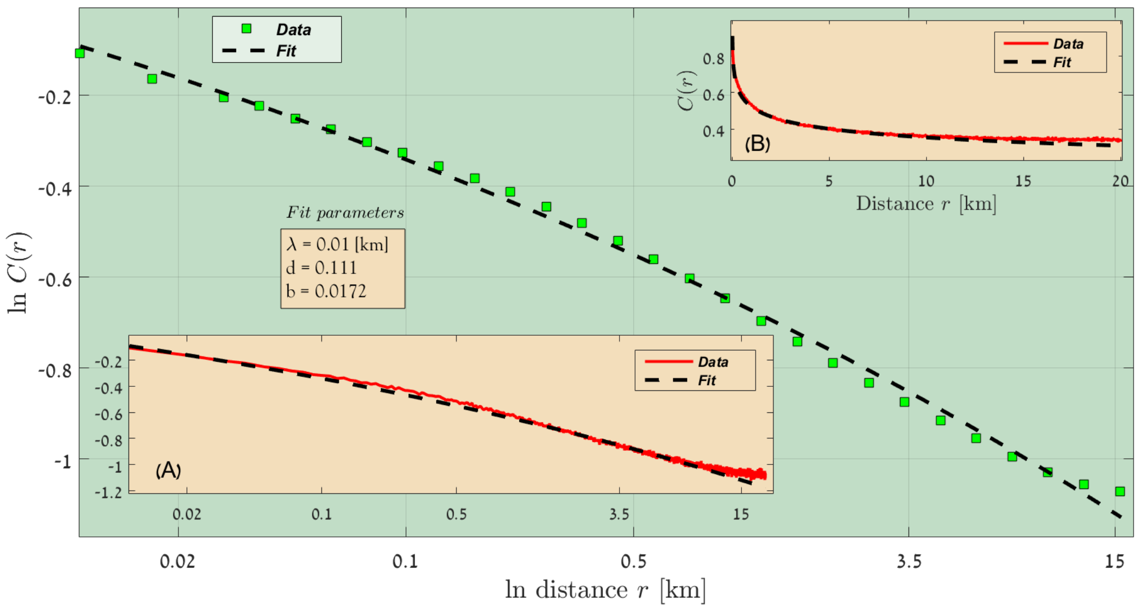


FIG. 62: Australia, Rainfall lines 300 - 350 [mm/year], 2016-2018.

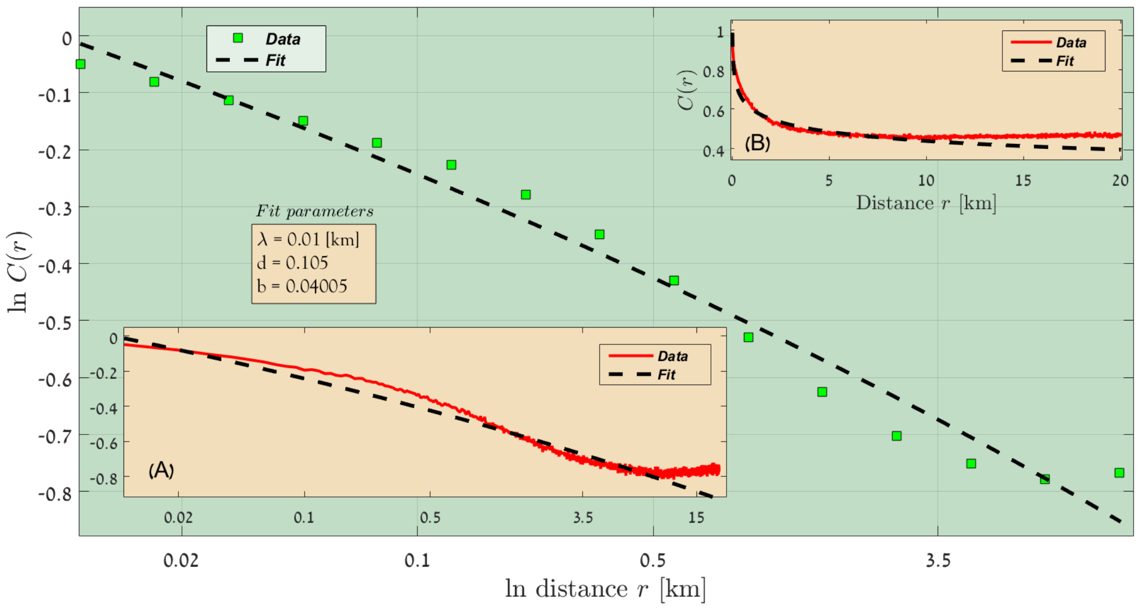


FIG. 63: Australia, Rainfall lines 350 - 400 [mm/year], 2016-2018.

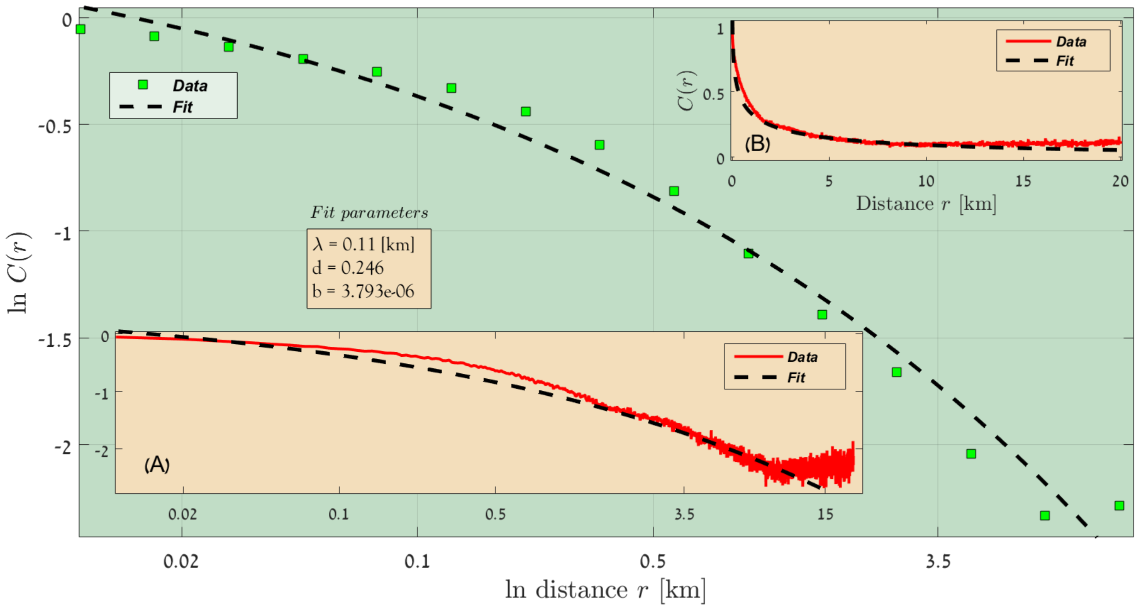


FIG. 64: Australia, Rainfall lines 400 - 450 [mm/year], 2016-2018.

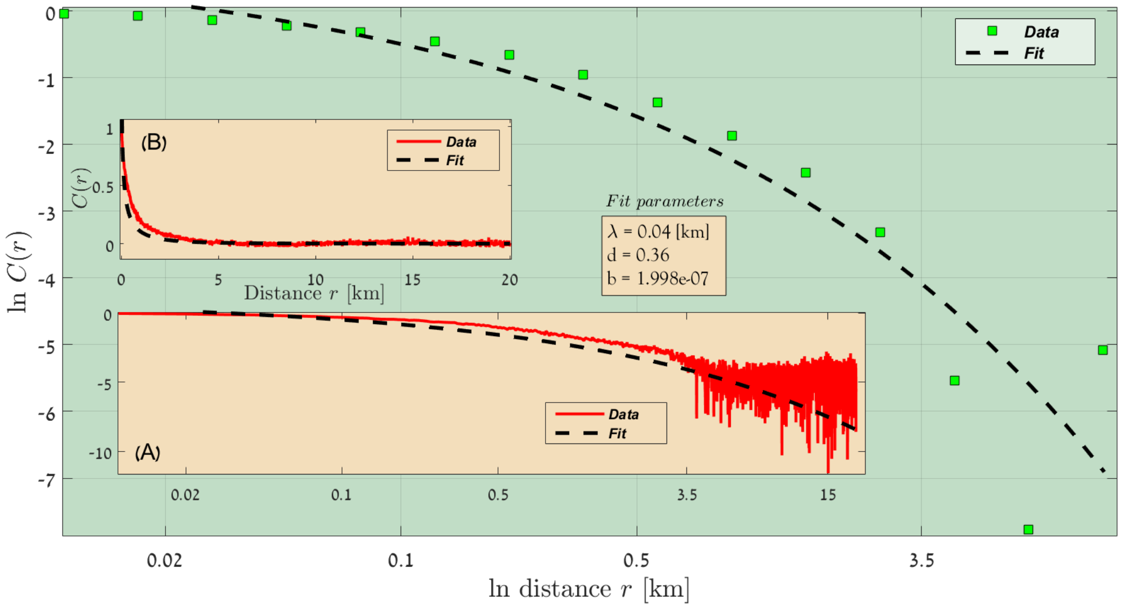


FIG. 65: Australia, Rainfall lines 450 - 500 [mm/year], 2016-2018.

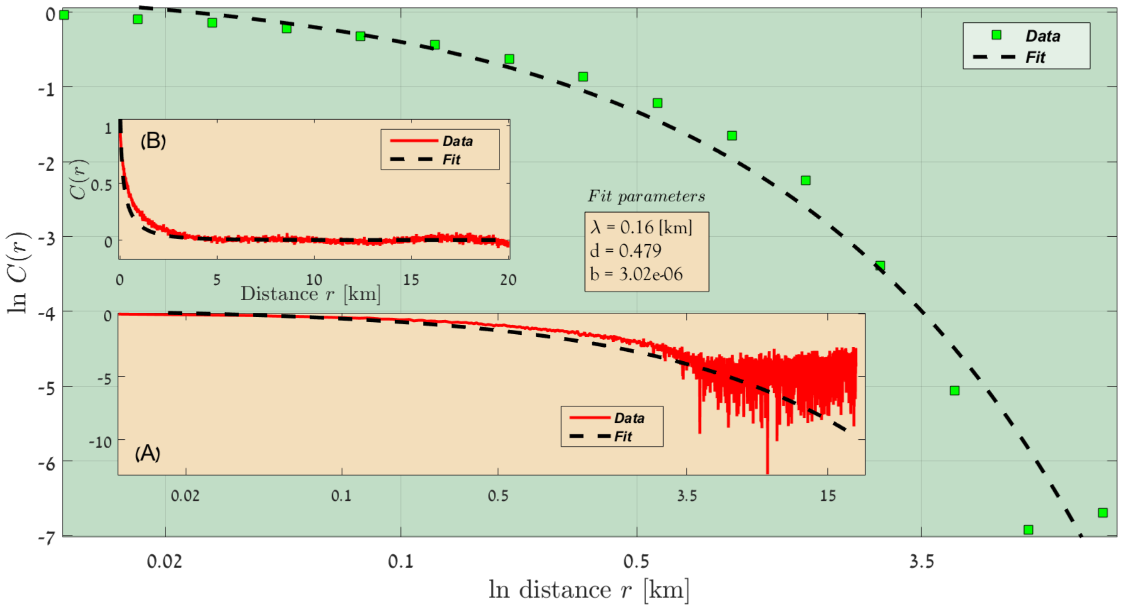


FIG. 66: Australia, Rainfall lines 500 - 550 [mm/year], 2016-2018.
